# Methylation-directed Acetylation of Histone H3 Regulates Developmental Sensitivity to Histone Deacetylase Inhibition

**DOI:** 10.1101/2020.07.22.215665

**Authors:** Li-Yao Huang, Duen-Wei Hsu, Catherine Pears

## Abstract

**Background:** Treatment of cells with hydroxamate-based lysine deacetylase inhibitors (KDACis) such as Trichostatin A (TSA) can induce biological effects such as differentiation or apoptosis of cancer cells, and a number of related compounds have been approved for clinical use. TSA treatment induces rapid initial acetylation of histone 3 (H3) proteins which are already modified by tri-methylation on lysine 4 (H3K4me3) while acetylation of bulk histones, lacking this mark, is delayed. Sgf29, a subunit of the SAGA acetyltransferase complex, interacts with H3K4me3 via a tandem tudor domain (TTD) and has been proposed to target the acetyltransferase activity to H3K4me3. However the importance of acetylation of this pool of H3 in the biological consequences of KDACi treatment is not known.

**Results:** We investigated the role of H3K4me3-directed acetylation in the mechanism of action of TSA on inhibiting development of the eukaryotic social amoeba *Dictyostelium discoideum*. Loss of H3K4me3 in strains with mutations in the gene encoding Set1 or the histone proteins confers resistance to TSA-induced inhibition of development and delays accumulation of histone acetylation on H3K9 and K14. A candidate orthologue of Sgf29 in *Dictyostelium* has been identified which specifically recognizes the H3K4me3 modification via its tandem Tudor domain (TTD). Disruption of the gene encoding Sgf29 delayed accumulation of H3K9Ac, abolished targeted H3K4me3-directed H3Ac and led to developmental resistance to TSA, which is dependent on a functional TTD. TSA resistance also results from overexpression of Sgf29.

**Conclusion:** Preferential acetylation of H3K4me3 histones, regulated by Sgf29 via its TTD, is important in developmental sensitivity to TSA. Levels of H3K4me3 or Sgf29 will provide useful biomarkers for sensitivity to this class of chemotherapeutic drug.

## Background

Specific post-translational modifications of histone proteins are associated with genes that are either active or repressed. For example, methylation of histone H3 on lysine 4 (H3K4) is a hallmark of genes able to be transcribed from yeast to humans, with the single lysine modified by the addition of up to three methyl groups. Within active genes, genome wide chromatin immunoprecipitation analysis has revealed that these modifications are region-specific. For example, H3K4me1 is associated with enhancer regions, while H3K4me2 and H3K4me3 are enriched at the promoter and regions proximal to the transcription start site (TSS) of actively transcribed genes[1]. Methylation of H3K4 is catalysed by lysine methyltransferases (KMTs) containing a characteristic SET domain[2]. In *Saccharomyces cerevisiae*, Set1 is the only methyltransferase catalysing *mono*, *di*- and *tri*-methylation of H3K4. The number of Set1 homologues expands in higher eukaryotes with three and six family members in *Drosophila melanogaster* and *Homo sapiens*, respectively, with some able to carry out *mono*, *di*- and *tri*-methylation of H3K4 while the others harbour only *mono*-to *di*-methylation ability[3].

Another modification associated with active genes is acetylation of multiple lysine residues in the N terminal tails of histones H3 and H4. For example, acetylation of H3K9 and H3K14 are often found together with H3K4me2/3 at the TSS region of actively transcribed genes[4]. H3K9 and H3K14 are substrates of members of the Gcn5 *N*-acetyltransferases (GNATs), MYST (Morf, Ybf2, Sas2 and Tip60) and p300/CBP acetyltransferases[5–7]. These histone modifications can be removed by lysine demethylases (KDMs) and deacetylases (KDACs) to allow dynamic turnover and resetting of modifications associated with a particular gene, without histone replacement.

Aberrant levels or locations of histone marks are associated with diseases such as cancer[8]. Mutations of genes encoding histone modifiers have been identified in numerous forms of cancer including mutation of KDMs and KDACs[9]. A classic example is the rearrangement of the MLL family of methyltransferases in acute lymphoblastic leukaemia and acute myeloid leukaemia[10], and a high number of somatic mutations of *MLL2* has been reported in patients with follicular lymphoma[11]. Therefore, histone modifiers are an attractive target for chemotherapy. Though development of small molecule inhibitors against KMT and KDM is underway, currently there are no clinically approved KMT or KDM inhibitors used to treat cancer. On the contrary, the development of KDAC inhibitors (KDACis) has proven success. Among six U.S. FDA-approved drugs against targets modifying chromatin structure, four of them are KDACis, namely Suberoylanilide hydroxamic acid (SAHA), Belinostat, Panobinostat and Romidepsin[9]. The hydroxamate KDACi Trichostatin A (TSA) is closely related to SAHA and often used experimentally. *In vitro* these drugs exert their effect by inducing either apoptosis or differentiation of cancer cells at concentrations that do not induce these cellular changes in non-cancer cells[12].

In humans 18 genes encoding KDACs have been assigned to four different classes based on their domain structure and sequence similarity. Class I, II, and IV are Zn^2+^-dependent while Class III KDACs are NAD^+^-dependent[13]. All four U.S. FDA-approved KDACis target Zn^2+^-dependent KDACs with different specificities[14]. Notably, SAHA, Belinostat and Panobinostat are all derivatives of hydroxamic acids. SAHA is approved for treatment of cutaneous T-cell lymphoma, Belinostat for peripheral T-cell lymphoma and Panobinostat for multiple myeloma. However, the response rate for patients diagnosed with these is around 30% and there are further problems with drug resistance emerging during treatment[15–17]. Mechanisms leading to resistance include elevated transporter activity, suppression of apoptosis, and escape from damage by reactive oxygen species[18]. Therefore, it is crucial to gain deeper understanding of the mode of action of hydroxamic acids in order to identify which cancers will respond to KDACi and to identify the mechanisms of inherent or induced resistance to improve the efficacy of these drugs.

KDACis have been proposed to exert their effects by inducing hyperacetylation on histone H3 and H4, resulting in alteration of gene expression and subsequent phenotypic changes[19]. Although acetylation is associated with upregulation of transcription of all genes, transcriptomic studies have found that treatment of cells with KDACis only changes the expression of a small portion of protein-coding genes (< 10%) with similar numbers both up- and down-regulated[20, 21]. Histone hyperacetylation induced by KDACi is not equally distributed across the whole genome so although KDACi treatment induces global histone hyperacetylation, some genomic regions are not acetylated and that is related to down-regulated genes[22]. Up-regulation of BIM, a pro-apoptotic protein in the Bcl-2 family, by Panobinostat in triple negative breast cancer cell lines was found coupled with induced levels of H3K27ac at two BIM enhancers[23].

There is another layer of complexity in KDACi-induced histone hyperacetylation. TSA- and SAHA-induced global H3K9/14Ac in human aortic endothelial cells is highly associated with up-regulated genes and co-localizes with H3K4me3[24]. Biochemical evidence suggests that KDACi-induced early hyperacetylation preferentially occurs on histones that are already carrying the H3K4me3 modification[25]. This reveals that H3K4me3 histones are normally subjected to rapid turnover of acetylation and so are the first histones to become acetylated upon TSA treatment. This dynamic acetylation of H3K4me3 histones, targeted to H3K9 and H3K14, is conserved across evolution in human, mouse and *Drosophila* cells[26]. However, it is unclear if this pool of histones subjected to H3K4me3-directed acetylation is a relevant target for the long-term biological effects of hydroxamates.

Unravelling the link between K3K4me3, histone acetylation and the biological effects of KDACi is hampered in higher eukaryotes by their complexity as a large number of enzymes are involved in adding and removing these modifications. We have circumvented this by investigating the role of dynamic acetylation in the mode of action of TSA using *Dictyostelium discoideum*. This is a haploid, single celled organism when feeding but undergoes multicellular development on starvation. *Dictyostelium* has many histone modifications conserved with higher eukaryotes but with a more limited numbers of proteins responsible for their deposition and removal[27, 28]. Notably, like yeast there is a single Set1 protein responsible for all detectable H3K4 methylation[29]. Also, unlike mammalian genomes which contain multiple copies of genes encoding most histones, *Dictyostelium* has single copies of the genes encoding two major histone H3 variants, H3a and H3b, facilitating mutation of the endogenous genes to probe the role of histone modifications. We have shown previously that in *Dictyostelium* H3K4me3-directed H3 acetylation (H3Ac) is conserved with higher eukaryotes[30]. Here we show that loss of methylation on H3K4, either due to deletion of *Set1* or mutation of the endogenous H3K4 genes, confers resistance to hydroxamate (TSA)-induced developmental inhibition and changes in histone acetylation profile. We identify a *Dictyostelium* homologue of Sgf29, a reader of H3K4me2/3 present in the Gcn5 HAT complex from yeast to humans and show that disruption of the gene encoding Sgf29 leads to loss of the rapid acetylation of H3K4me3 histones and also confers TSA resistance. We further demonstrate the resistance to TSA can also be achieved by overexpressing Sgf29. This highlights the dynamically acetylated pool of H3K4me3 histones as an important target for the biological consequences of TSA treatment and suggests that the methylation status of H3K4 as well as the levels of Sgf29, will influence cellular sensitivity to hydroxamates providing potential novel biomarkers for tumour cell sensitivity.

## Results

### Dictyostelium development is inhibited by exposure to TSA in early stages

In *Dictyostelium* the developmental cycle is triggered by starvation and involves initial aggregation of individual amoebae to form a multicellular structure. This goes on to generate a fruiting body after 24 hours, consisting of a spore head raised from the surface by a stalk consisting of dead vacuolated cells. TSA inhibits *Dictyostelium* cellular KDAC activity and development[31]. This therefore provides a quick and reliable system to assess the biological consequences of treatment with hydroxamates in a system with complexity including differentiation of different cell types, cell movement and programmed cell death.

To confirm the dose dependence and stage of inhibition, *Dictyostelium* Ax2 cells were developed by starvation on buffered agar containing increasing concentrations of TSA, and analysed throughout the life cycle until vehicle-treated controls had mostly culminated to form fruiting bodies (Fig. 1a). In the presence of all concentrations of TSA, aggregates formed at 8 hours, a time similar to the DMSO treated controls, but a delay in culmination was apparent at concentrations of 1 μM and above. From 2 μM TSA, most of the structures observed after 24 h of development were at still at the earlier aggregate stage. Indeed, the percentage of structures past the aggregation stage (“late structures”) decreased significantly at 2 μM and 4 μM compared to control cells (Fig. 1a). The result suggests that transition from aggregation to culmination was inhibited by TSA.

**Figure. 1.**
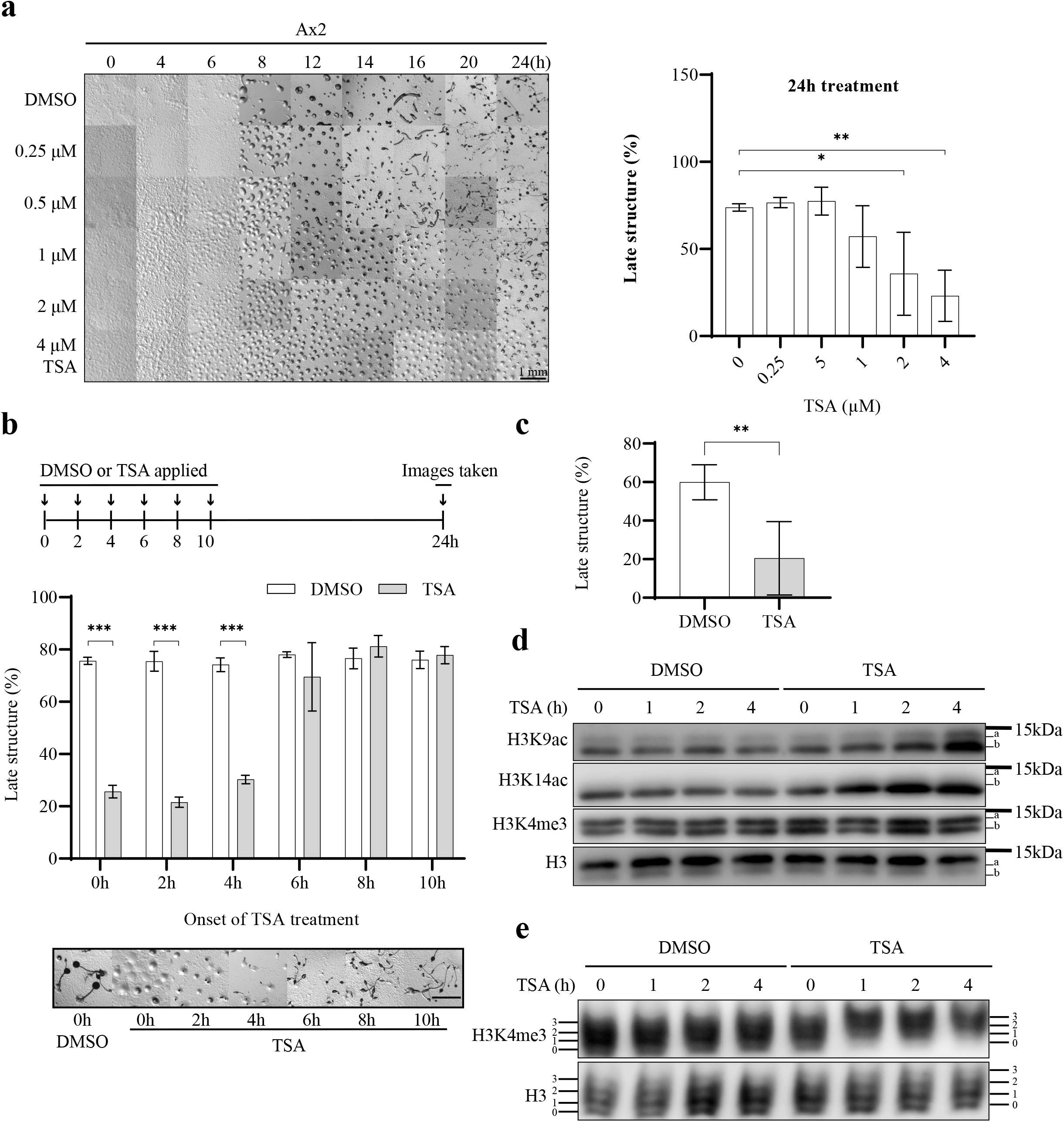
Early exposure to TSA induces H3K4me3-directed H3 hyperacetylation and inhibits later development of *Dictyostelium*. **a** TSA inhibits the development of *Dictyostelium*. Left panel, exponential Ax2 cells were collected, washed twice with KK2 and allowed to develop on 1.5% (w/v) agar (1.5 × 10^6^ cells/cm^2^) containing increasing concentrations of TSA (0 – 4 μM). Images were taken at indicated time and representative images of three repeats presented. Right panel, Percentage of late structures (beyond aggregation stage) was assessed at 24 hours (average of n=3 ± SD). Statistical significance was calculated using two-way ANOVA with Tukey test. **b** Early exposure of cells to TSA is necessary for developmental inhibition. Upper panel, schematic illustration of the experimental design. The development assay was performed as described in a, but TSA (4 μM) was added at the indicated time during development and pictures taken after 24 hours. Middle panel, quantification of late structures of three biological repeats. Bottom panel, representative images. **c.** TSA-treatment for four hours inhibits late development. Cells were washed using KK2 and incubated with DMSO or 4 μM TSA for four hours in KK2 in shaking suspension at 4 × 10^6^ cells/ml and subsequently transferred onto agar for development for another 20 hours before images were taken and percentage of late structures was calculated. Data in b and c is the average ± SD of three independent repeats and statistical significance calculated using Student’s paired T-test**. d** TSA-induced changes in H3K9Ac, H3bK14Ac. Cells were washed in KK2 and developed in the presence of 4 μM TSA or DMSO vehicle control for 0 - 4 hours at 4 × 10^6^ cells/ml in KK2 before harvesting. Acid-extracts were resolved by 18% SDS-PAGE and immunoblotted using specific antibodies as indicated. H3 was used as a loading control. **e** TSA induces H3K4me3-directed acetylation. Cells were treated and harvested as described in d. Acid-extracts were resolved on 20% Acid-urea gels and immunoblotted using anti-H3K4me3 and H3 antibodies. *, *p* < 0.05; **, *p* < 0.01; ***, *p* < 0.001.

To address whether TSA is having an effect post-aggregation or whether the late block reflects the effects of TSA earlier in development, cells were allowed to develop for 24 hours on agar and DMSO or 4 μM TSA were added at different timepoints (Fig. 1b). All control DMSO-treated samples formed fruiting bodies at 24 hours (Fig. 1b). TSA-induced developmental inhibition was observed in samples where TSA was added after 0, 2 and 4 hours of development but not in samples exposed to TSA after 6 hours. This suggests that the block in late developmental is a consequence of earlier exposure to TSA. To confirm this, Ax2 cells were exposed to 4 μM TSA (or DMSO control) by developing in shaking suspension for the first four hours, then washed to remove the TSA before being allowed to develop on agar for another 20 hours in the absence of TSA. While 60% of the total structures passed the aggregate stage when cells were pre-treated with DMSO, the number dropped significantly to 20% for TSA-treated cells (Fig. 1c). This confirms that treating developing cells with TSA for the first four hours of development is sufficient to reduce the percentage of late structures while also suggesting that at least some of the molecular event(s) leading to developmental inhibition by TSA happen during the first fours of development.

### TSA induces H3K9/K14 acetylation and preferential acetylation of H3K4me3 in developing cells

To understand the temporal change of histone acetylation during the first four hours of development, exponentially growing Ax2 cells were starved in shaking suspension in the presence or absence of 4 μM TSA for up to 4h. Levels of H3K9Ac, H3bK14Ac and H3K4me3 were detected by western-blot, using modification-specific antisera which have been verified for specificity in *Dictyostelium*[25, 26, 30]. The two main histone H3 variants expressed in *Dicytostelium,* H3a and H3b, can be separated on high resolution gels as H3a contains three extra amino acids[30] allowing modification of both variants to be assessed. In the absence of TSA, H3K9Ac and H3bK14Ac were apparent in developing cells, mainly on H3b, and the levels did not markedly change during the first four hours of development. However, when developed in the presence of TSA, H3K9Ac and H3bK14Ac levels increased from 1 hour of TSA treatment (Fig. 1d). In contrast, the levels of H3K4me3 remained unchanged in control and TSA-treated cells across the four hours. Thus, developing cells exposed to TSA in early development accumulate H3K9Ac and H3bK14Ac, and the change is not due to development.

In order to investigate the role of preferential acetylation of H3K4me3 histones in the mode of action of TSA, it is necessary to determine if this increase in acetylation is preferentially targeted to H3K4me3 histones in developing cells as previously reported for growing *Dictyostelium* cells[30]. Therefore, histone-enriched samples were resolved using acid-urea (AU) gel electrophoresis which separates proteins by charge as well as size. Methylation does not alter the charge, but acetylation of a single lysine reduces the positive charge by one, leading to a ladder of histones (level 0 – 3) detectable by western-blot corresponding to different acetylation states, with the least acetylated running further in the gel. The acetylation state of both total histone H3 and H3K4me3 were detected by western-blot. Consistent with the SDS-PAGE western-blot analysis (Fig. 1d), neither total histone H3 or H3K4me3 histones showed any increase in acetylation during the first 4 hours of development (Fig. 1e, lane 1 - 4). In contrast, cells developed in the presence of 4 μM TSA showed a shift of bands in H3K4me3 ladder. The major bands of the H3K4me3 ladder shifted from level 1 and 2 to level 2 and 3 within 1 hour, while bulk histone H3 remained the same across the course of treatment (Fig. 1e lane 5 – 8). Taken together, TSA can induce rapid acetylation of histone H3 in developing cells and H3K4me3 histones are preferentially acetylated, confirming H3K4me3-directed H3 acetylation (H3Ac) during early development.

### set1^−^ cells lacking H3K4me are resistant to TSA treatment during development

To understand whether H3K4me3 histones are a biologically relevant target for TSA, firstly the developmental sensitivity to TSA treatment of cells lacking H3K4 methylation was determined. In *Dictyostelium*, Set1 is the sole enzyme responsible for *mono*-, *di*- and *tri*-methylation of H3K4[29]. Therefore, Ax2 and *set1*^−^cells were developed in the presence of 0 – 4 μM TSA treatment (Fig. 2a). As before, development of Ax2 cells was markedly inhibited, showing a significant decrease of percentage of late structures starting from 1 μM TSA. However, *set1*^−^ cells successfully formed fruiting bodies even at the highest concentration of TSA tested, and the percentage of late structures remained close to untreated cells (Fig. 2a). The kinetics of histone H3 acetylation was also examined (Fig. 2b). In contrast to the rapid accumulation of H3 acetylation in Ax2 cells, *set1*^−^ cells failed to accumulate a significant increase in levels of H3K9Ac and H3bK14Ac upon TSA treatment during the first four hours of development. The H3K4me3 level did not change significantly in Ax2 cells and was not detectable in *set1*^−^ cells as expected. Therefore, the effect of TSA on histone acetylation of these residues is delayed in *set1*^−^ cells correlating with their resistance to TSA-induced developmental inhibition.

**Fig 2.**
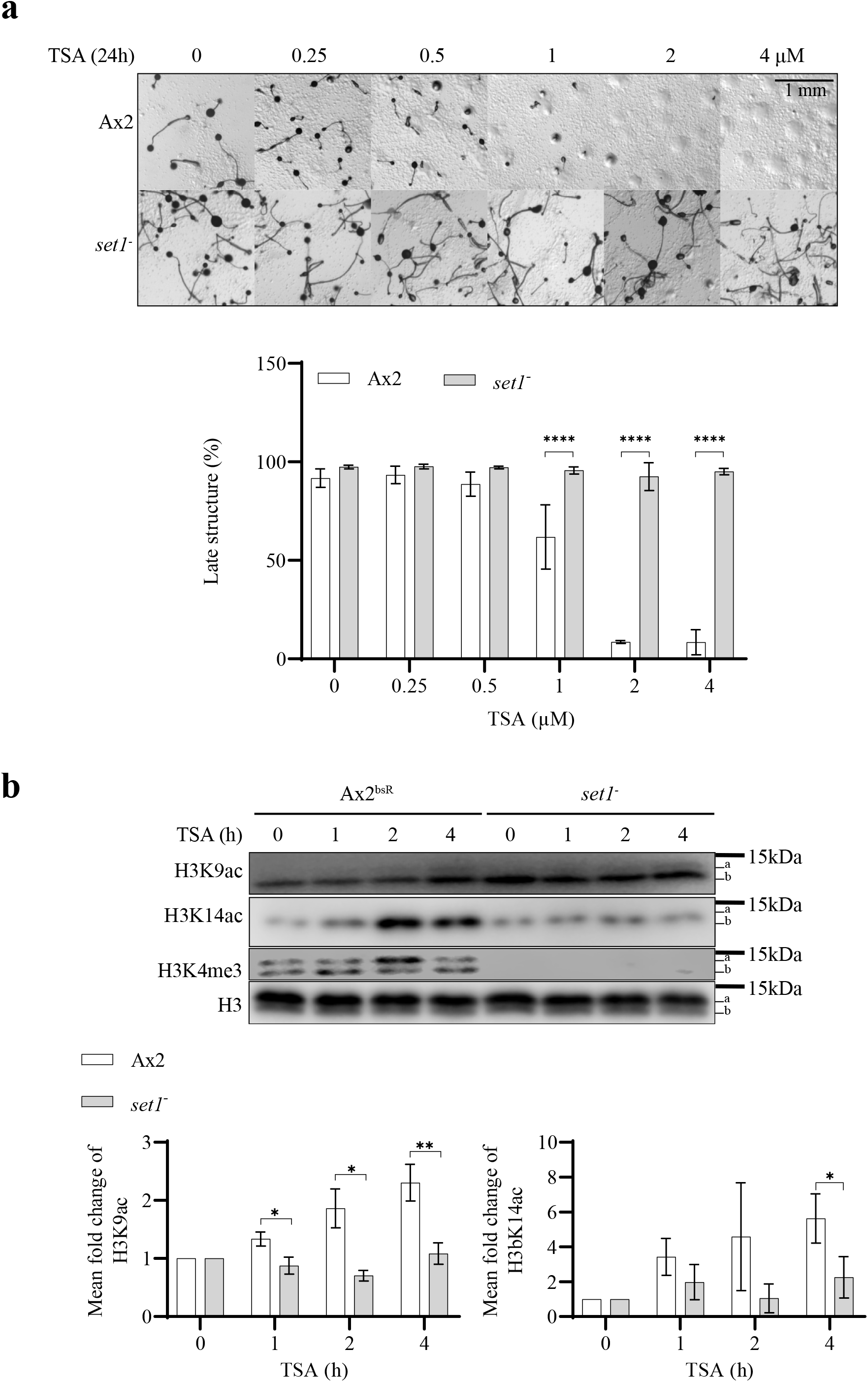
Loss of Set1 delays H3 acetylation and confers developmental resistance to TSA. **a** *set1*^−^ cells are resistant to TSA during development. Ax2 and *set1*^−^ cells were washed with KK2 and allowed to develop on solid agar with increasing concentrations of TSA for 24 hours as described in Fig. 1a. Images were taken at 24 hours and representative images of three repeats were presented. Percentage of late structures was calculated and shown as average ± SD. Statistical significance was calculated using two-way ANOVA with Tukey test. **b** TSA-induced change of H3 acetylation is delayed in *set1*^−^ cells. Cells were developed in the presence of 4 μM TSA for 0 - 4 hours in KK2 before harvesting for acid-extraction as described in Figure 1. Acid-extracts were resolved by 18% SDS-PAGE and immunoblotted using specific antibodies as indicated. Quantification of H3K9Ac and H3bK14Ac relative to total H3 was calculated from three biological repeats and shown as mean ± SD. Value of 0h was set as 1. S tatistical significance was calculated using Student’s paired T-test. *, *p* < 0.05; **, *p* < 0.01; ****, *p* < 0.0001.

### H3K4A cells are resistant to TSA during development

It is known that methyltransferases containing SET domains can have substrates other than histones[32]. Therefore, in order to determine whether the phenotypes of *set1*^−^ cells are due to loss of H3K4 methylation, we utilized two strains in which the endogenous single copy histone H3 genes had been replaced with mutated versions to introduce an H3K4A mutation on either of the H3 variants, H3a or H3b[30]. It has previously been reported that H3aK4A cells have no detectable H3K4me3 on H3a or H3b while H3bK4A cells maintain detectable H3K4me3 on H3a but not H3b[30]. H3aK4K cells (a gene replacement including insertion of the Blasticidin resistance cassette but without the introduced mutation) served as a control.

Similar to parental Ax2 cells, development of H3aK4K cells was markedly inhibited at concentrations of TSA above 1 μM after 24 hours of treatment. In contrast, H3aK4A and H3bK4A cells were able to develop beyond the aggregate stage (Additional File 1: Figure S1). However, unlike *set1*^−^ cells which formed mature fruiting bodies at concentrations of up to 4 uM TSA within 24 hours (Fig. 2a), H3aK4A and H3bK4A cells failed to form mature fruiting bodies after 24h hours of TSA treatment. However, after 48h, both H3aK4A and H3bK4A exhibited markedly higher percentage of late structures at concentrations above 1 μM TSA when compared to H3aK4K cells (Fig. 3a). This suggests that loss of methylation on H3K4 is at least a contributor to the resistance phenotype. Furthermore, loss of H3K4me3 on H3b alone is sufficient to confer TSA resistance during development. Thus, we further analyzed the change of H3 acetylation in developing H3bK4A cells exposed to 4 μM TSA (Fig. 3b).

**Fig 3.**
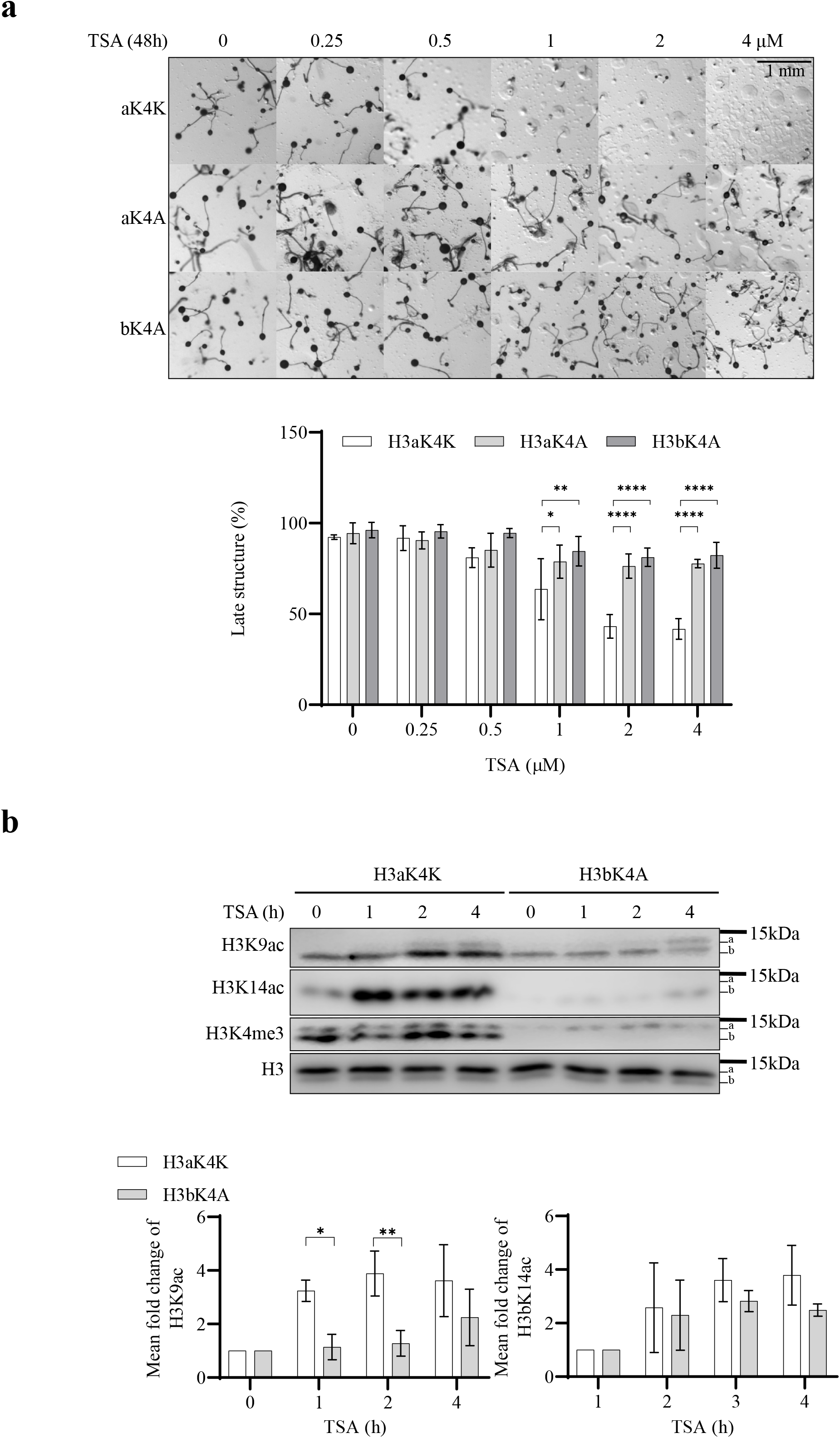
Loss of H3K4me3 delays H3 acetylation and confers developmental resistance to TSA. **a** H3aK4A and H3bK4A cells are resistant to TSA during development. H3aK4A and H3bK4A and control H3aK4K cells were washed with KK2 and allowed to develop on solid agar with increasing concentration of TSA for 24 hours as described in Fig. 1a. Images were taken at 24 hours and representative images of three repeats were presented. Percentage of late structures was calculated and shown as average ± SD. Statistical significance was calculated using two-way ANOVA with Tukey test. **b** TSA-induced change of H3 acetylation is delayed in H3bK4A cells. Cells were washed with KK2 and developed in the presence or absence of 4 μM TSA for 0 - 4 hours in KK2 before harvesting for acid-extraction. Acid-extracts were resolved in 18% SDS-PAGE and immunoblotted using specific antibodies as indicated. Quantification of H3K9Ac and H3bK14Ac relative to total H3 was calculated from three biological repeats and shown as mean ± SD. Value of 0h was set as 1. S tatistical significance was calculated using Student’s paired T-test. *, *p* < 0.05; **, *p* < 0.01; ****, *p* < 0.0001.

Similar to parental Ax2 cells, the level of H3K9Ac in H3aK4K cells was significantly higher than H3bK4A cells after 1h of development in the presence of TSA (Fig. 3b). H3bK4A showed no significant accumulation of H3K9Ac on H3b. However, on H3a (which is still methylated on K4) there was a slight increase in H3K9Ac at the 4h timepoint (Fig. 3b). In both H3aK4K and H3bK4A cells, H3bK14Ac accumulation during development in the presence of TSA was observed. However, the overall signal was markedly weaker in H3bK4A cells compared to H3aK4K cells. The H3K4me3 level did not change significantly in H3aK4K cells and the signal on H3b was not detectable in H3bK4A cells as expected. Thus, loss of modification on H3bK4 resulted in loss of rapid increase in levels of H3bK9ac in the presence of TSA. In *Dictyostelium*, the only known modification on H3K4 is methylation[28]. This suggests that methylation of H3K4 is necessary for rapid TSA-induced accumulation of H3K9Ac.

### Loss of Sgf29 causes delayed accumulation of H3 acetylation upon TSA treatment

In *S. cerevisiae*, Sgf29 is a subunit of the HAT module of the histone acetyltransferase SAGA (Spt-Ada-Gcn5-acetyltransferase) complex. Sgf29 contains a tandem Tudor domain (TTD) that preferentially binds to H3K4 *di*- and *tri*-methylated histones *in vitro* and has been proposed to link H3K4me2/3 and histone acetylation[33]. In order to directly test whether the H3K4me3-directed H3Ac is involved in TSA resistance by generating a strain in which H3K4me3 is conserved but its preferential acetylation is lost, we identified a candidate Sgf29 ortholog in *Dictyostelium*. The protein sequences of Sgf29 from various species share low identity in the *N*-terminal region but have a relatively well-conserved TTD sequence at the *C*-terminus[33]. Therefore, the protein sequences of the TTD of budding yeast, fission yeast and humans were used in three independent BLAST searches against predicted *Dictyostelium* proteins (http://dictybase.org/). All three searches resulted in homology with only one candidate protein of 411 amino acids encoded by the gene DDB_G0272054, which, like Sgf29 from other organisms, is predicted to have a TTD at the *C*-terminus (Fig. 4a). In addition, three residues (F359, Y366 and F387) in the TTD required for interaction with *tri*-methylated lysine H3K4 in yeast[33] are conserved, though in *Dictyostelium* the first aromatic residue is a tyrosine rather than a phenylalanine. A structure alignment (Additional File 2: Figure S2) is consistent with the TTD of the *Dictyostelium* protein being structurally conserved with that of yeast and humans[33, 34].

**Fig 4.**
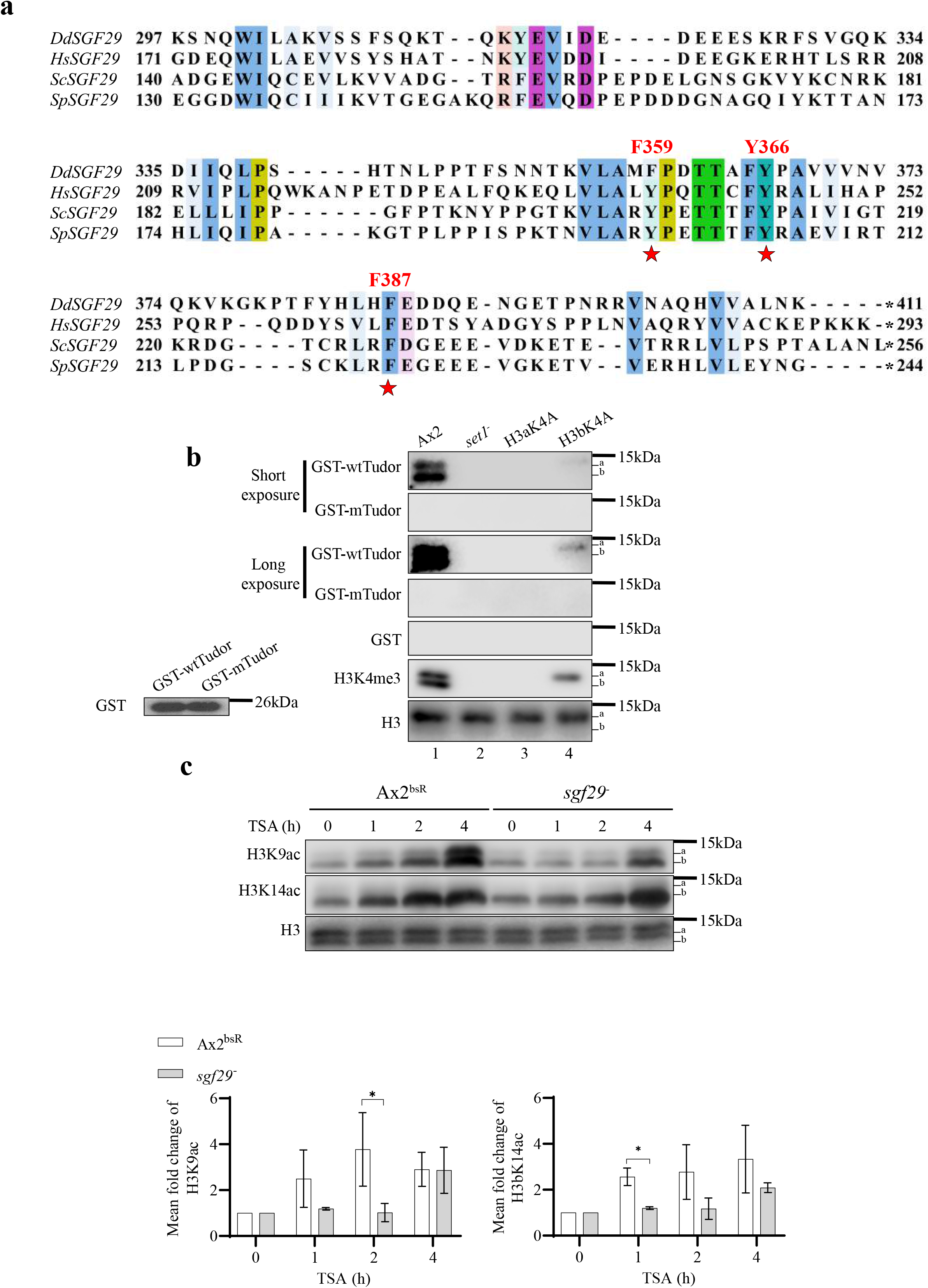
TTD of Sgf29 recognizes H3K4me3 while deletion of Sgf29 delays TSA-induced rapid H3 hyperacetylation. **a** A candidate *Dictyostelium* orthologue of Sgf29 (DdSgf29, 411 a.a.) contains conserved tandem Tudor domain (TTD). Protein sequence alignment of the TTD of Sgf29 from budding yeast (Sc), fission yeast (Sp) and human (Hs) together with DdSgf29. Conserved binding residues targeting *tri*-methylated lysine are labelled with red asterisks and their positions relative to the first amino acid are shown. *, stop codon. Clustal × Colour Scheme is used to indicate the conservation of amino acids. **b**. TTD interacts with H3K4me3 histones. Wildtype or F359A/Y366A TTD were expressed and purified from *E.coli* as GST-fusion proteins. Acid extracts were prepared from exponentially growing Ax2, *set1^−^*, H3aK4A and H3bK4A cells and resolved by SDS-PAGE, transferred to a PVDF membrane and incubate with one of the fusion proteins. The interaction between GST-fusion protein and histone H3 is detected using an anti-GST monoclonal antibody. GST alone was used as a negative control. Equal amount of purified recombinant proteins was used as indicated by the lower left blot. Specific antibodies were used to detect H3K4me3 and histone H3. Representative blot (*n* = 3). **c** TSA-induced change of H3 acetylation is delayed in *sgf29*^−^ cells. Control Ax2^bsR^ cells (with random integration of the gene disruption vector) and *sgf29*^−^ cells were washed with KK2 and developed in the presence or absence of 4 μM TSA for 0 - 4 hours in KK2 before harvesting for acid-extraction. Acid-extracts were resolved in 18% SDS-PAGE and immunoblotted using specific antibodies as indicated. Quantification of H3K9Ac and H3bK14Ac relative to total H3 was calculated from three biological repeats and shown as mean ± SD. Value at 0h was set as 1. Statistical significance was calculated using Student’s paired T-test. *, *p* < 0.05.

The ability of the TTD of this *Dictyostelium* protein to interact with H3K4-trimethylated histones was tested using a far-western-blot strategy. Either wildtype TTD or a version containing two point mutations (F359A and Y366A) shown to disrupt binding to methylated H3K4 in the yeast version[33] were expressed in *E. coli* and purified as GST fusion proteins (Additional File 3: Figure S3). Equal amounts of the purified protein were used to detect interaction with histone-enriched acid extracts (Fig. 4b). Wildtype GST-TTD interacts with a band of the predicted molecular weight of histone H3 from Ax2 and H3bK4A cells but not *set1*^−^ or H3aK4A cells which both lack the H3K4me3 modification (Fig 4b). Notably, no detectable interaction with H3b was observed in H3aK4A cells, suggesting the binding affinity of GST-TTD to H3K4me3 is stronger than H3K4me1/2, which is still present in these cells[30]. In the contrast, GST-F359AY366ATTD failed to detectably interact with histone H3 from any of the cells. Together, this demonstrates that the TTD interacts with H3K4me3 histones *in vitro* so this protein will be referred so as DdSgf29.

*sgf29*-null strains were created (Additional File 4: Figure S4) and tested for histone acetylation in response to TSA during development. Similar to previous results, control Ax2^bsR^ cells accumulated H3K9Ac and H3bK14Ac starting from 1 hour of TSA treatment in the presence of TSA (Fig. 4c. In contrast, accumulation of H3K9Ac and H3bK14Ac in *sgf29*^−^ cells were first seen only after 4 hours of development in the presence of TSA (Fig. 4c). Therefore, disruption of *sgf29* in *Dictyostelium* causes delay of TSA-induced accumulation of H3 acetylation during development, a phenotype shared with *set1*^−^ and H3bK4A cells.

### Sgf29^−^ cells lack H3K4me3-directed H3Ac and are more resistant to TSA during development

To understand if the delayed accumulation of H3 acetylation is particularly targeted to H3K4me3 histones, acetylation of histones from *Sgf29*^−^ and control cells (Ax2^bsR^) were analysed by AU gel electrophoresis. As previously, more rapid acetylation of H3K4me3 than of total histone H3 was apparent in Ax2^bsR^ cells. The percentage of level 0 and 1 band of H3K4me3 significantly decreased at 2 hours from 22% and 37% to 9% and 30%, respectively. Concurrently, the percentage at levels 2 and 3 increased from 30% and 10% to 37% and 22%, respectively. Thus, the major H3K4me3 bands shifted from low acetylation states (level 0 and 1) to high acetylation states (level 2 and 3) within 2 hours of TSA treatment. However, any shift to higher acetylation states for bulk histone H3 was only observed after 4 hours, and the change was not significant (Fig 5a, lane 1 – 4). In *Sgf29*^−^ cells, this rapid acetylation of H3K4me3 was lost as the shift of H3K4me3 signal to levels 2 and 3 was only apparent at 4 hours together with the bulk histone H3. Moreover, the shift was only significant for H3 but not H3K4me3 histones (Fig 5a, lane 5 – 8). These findings were verified in two further independently derived *sgf29*^−^ clones. (Additional File 5: Figure S5a). Thus, *DdSgf29* is required for targeting preferential acetylation to H3K4me3 *in vivo* in *Dictyostelium* during development. Consistent with the importance of this acetylation, *sgf29*^−^ cells are resistant to TSA during development (Fig. 5b), verified in two further independent clones (Additional File 5: Figure S5b). Thus, disruption of the gene encoding Sgf29 not only results in delay of histone acetylation and loss of rapid H3K4me3-directed H3Ac, but also makes cells resistant to TSA-induced developmental inhibition (Fig. 4 and 5). These results support the hypothesis that Sgf29 mediates the crosstalk between H3K4me3 and histone acetylation, and that crosstalk is important for the mode of action of TSA.

**Fig 5.**
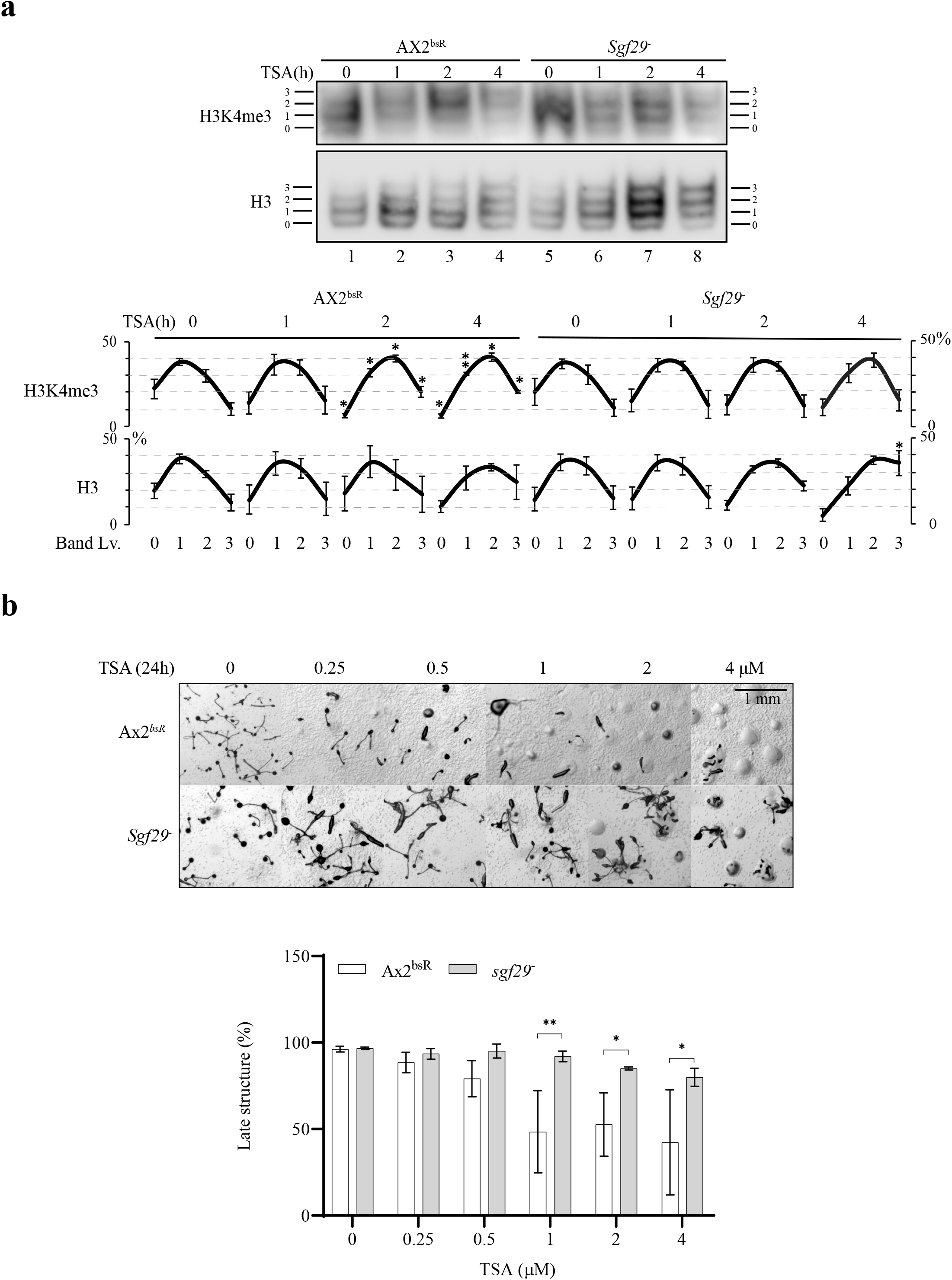
*sgf29*^−^ cells have lost rapid H3K4me3-directed H3 acetylation and are resistant to TSA during development. **a** H3K4me3-directed H3 acetylation is lost in *sgf29*^−^ cells. Acid extracts from Ax2^bsR^ and *sgf29*^−^ cells were prepared after 0, 1, 2 or 4 hours of development in the presence or absence of 4 μM TSA. Samples were resolved by 20% acid-urea electrophoresis and western blots were performed using anti-H3K4me3 and anti-H3 polyclonal antibodies. Positions of histone H3 with decreasing net positive charge (0 – 3) are shown on both sides of each blot. Representative blots (*n* = 3). Quantification of individual bands was shown as percentage of total intensity ± SD. Statistical significance was calculated by comparing to 0h using Student’s paired T-test. **b** *sgf29*^−^ cells are resistant to TSA during development. Cells were washed with KK2 and allowed to develop on solid agar with increasing concentrations of TSA for 24 hours as described in Fig. 1a. Images were taken at 24 hours and representative images of three repeats were presented. Percentage of late structures was calculated and shown as average ± SD. Statistical significance was calculated using two-way ANOVA with Tukey test. *, *p* < 0.05; **, *p* < 0.01.

### The role of Sgf29 in TSA sensitivity is dependent on the tandem Tudor domain

To confirm that the phenotypes seen above were due to loss of Sgf29 expression and to determine whether this was dependent on the ability of the protein to bind H3K4me3, constructs to drive expression of epitope tagged full length Sgf29-FLAG or Sgf29F359A/Y366A-FLAG from its endogenous promoter (containing sequences 831 bp upstream of the start codon) were generated (Fig. 6a). Single copy insertion required blasticidin as a selectable marker so the BsR cassette used to create the gene disruption in *sgf29*^−^ strain was first removed using the cre-lox system as previously described[35]. Expression of similar levels of Sgf29-FLAG or Sgf29F359A/Y366A-FLAG was confirmed using anti-FLAG-antibody (Fig. 6b). Expression of Sgf29 in *sgf29*^−^ cells re-sensitizes cells to TSA to a level similar to Ax2 cells during development (Fig. 6b). At concentrations above 2 μM TSA, the percentage of late structures of *sgf29*^−^ cells was significantly higher than cells re-expressing Sgf29. In the contrast, the number remained at 90% in cells re-expressing Sgf29F359A/Y366A at 2 μM TSA treatment but at 4 μM TSA dropped. significantly to 45% compared to *sgf29*^−^ cells (Fig.6B, lower panel). The result confirms that loss of Sgf29 is responsible for the TSA resistance in *sgf29*^−^ cells while also suggesting that the mutated Sgf29F359A/Y366A still has a partial function *in vivo*.

**Fig 6.**
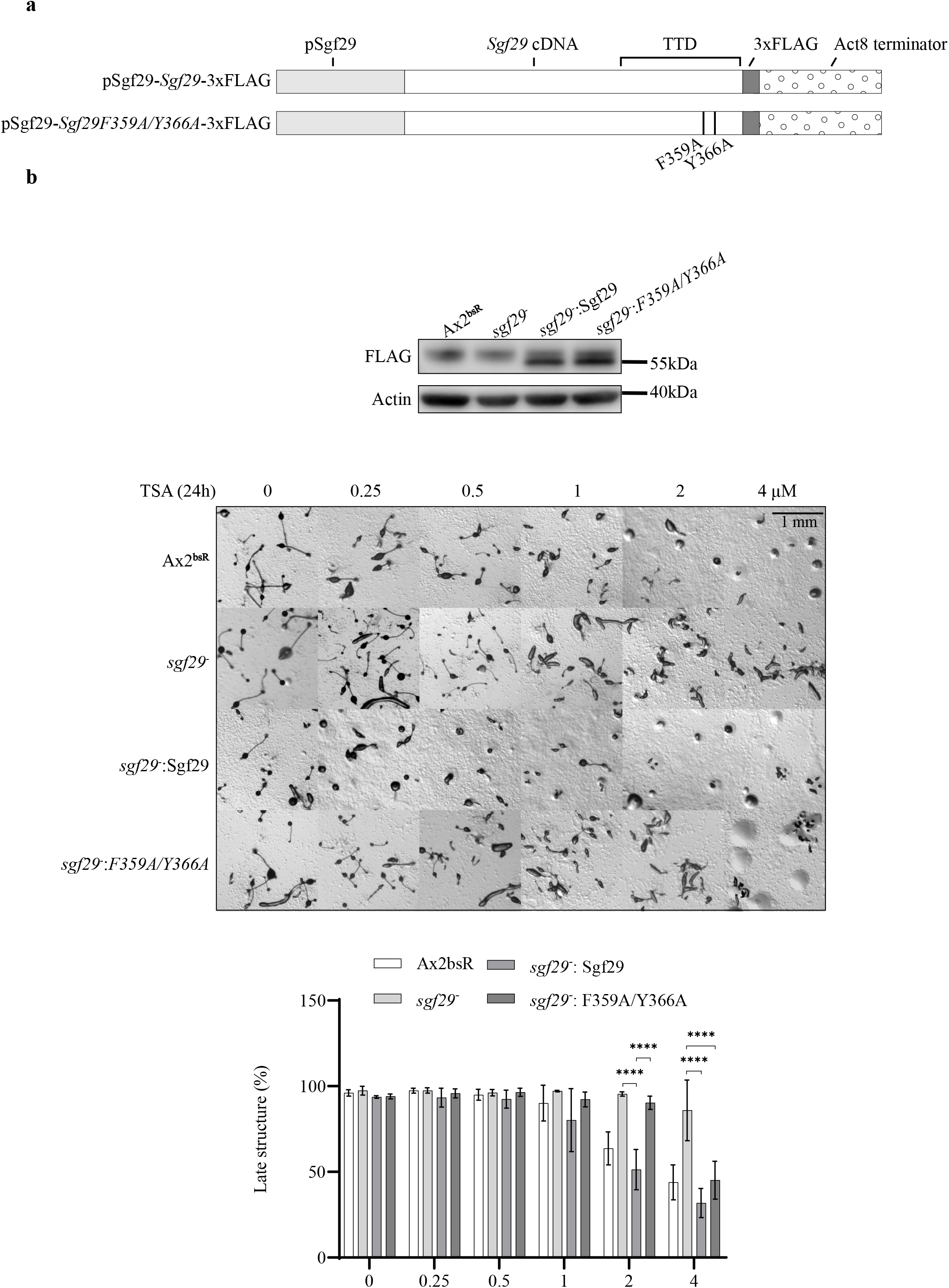
Cell sensitivity to TSA depends on functional TTD of Sgf29. **a** Schematic illustration of wildtype and F359A/Y366A Sgf29 constructs. The expression of *C*-terminally 3xFLAG-tagged protein is driven by the endogenous promoter of *sgf29* in an integrating vector **b** Expression of Sgf29-FLAG in *sgf29*^−^ cells re-sensitizes cells to TSA during development. Upper panel, immunoblots showing expression of 3xFLAG-tagged wildtype and F359A/Y366A Sgf29 in *sgf29*^−^ cells. Whole cell lysates from control cells, *sgf29*^−^ cells and *sgf29*^−^ cells expressing wildtype or F359A/Y366A Sgf29-FLAG was resolved by 12% SDS-PAGE and blotted using anti-FLAG antisera. Actin is used as a loading control. Lower panel, cells were washed with KK2 and allowed to develop on solid agar with increasing concentration of TSA for 24 hours as described in Fig. 1a. Images were taken at 24 hours and representative images of three repeats were presented. Percentage of late structures was calculated and shown as average ± SD. Statistical significance was calculated using two-way ANOVA with Tukey test. Statistical significance was calculated by comparing to 0h using one-way ANOVA with Dunnett test. ****, *p* < 0.0001.

### Overexpression of Sgf29 also leads to TSA resistance during development

As a number of tumour cells show increased levels of Sgf29 (COSMIC database COSU376, COSU377 and COSU414) we tested whether overexpression would also alter TSA sensitivity. Constructs to drive overexpression of full length Sgf29-FLAG or Sgf29F359A/Y366A-FLAG driven by the strong actin 15 promoter in an extrachromosomal, multicopy vector (Fig. 7a) were introduced into Ax2 cells and expression confirmed. (Fig 7b). The development of Ax2 cells overexpressing Sgf29 was markedly more resistant to TSA than control cells. In addition, Ax2 cells overexpressing Sgf29F359A/Y366A-FLAG exhibited mild resistance with the formation of late structures significantly inhibited down to 26% by 4 μM TSA compared to 7% in cells with empty vector and 86% in cells overexpressing Sgf29-FLAG (Fig. 7b). These results suggest that the level of Sgf29 expression defines the sensitivity of response to TSA as either its loss or overexpression leads to TSA resistance.

**Fig 7.**
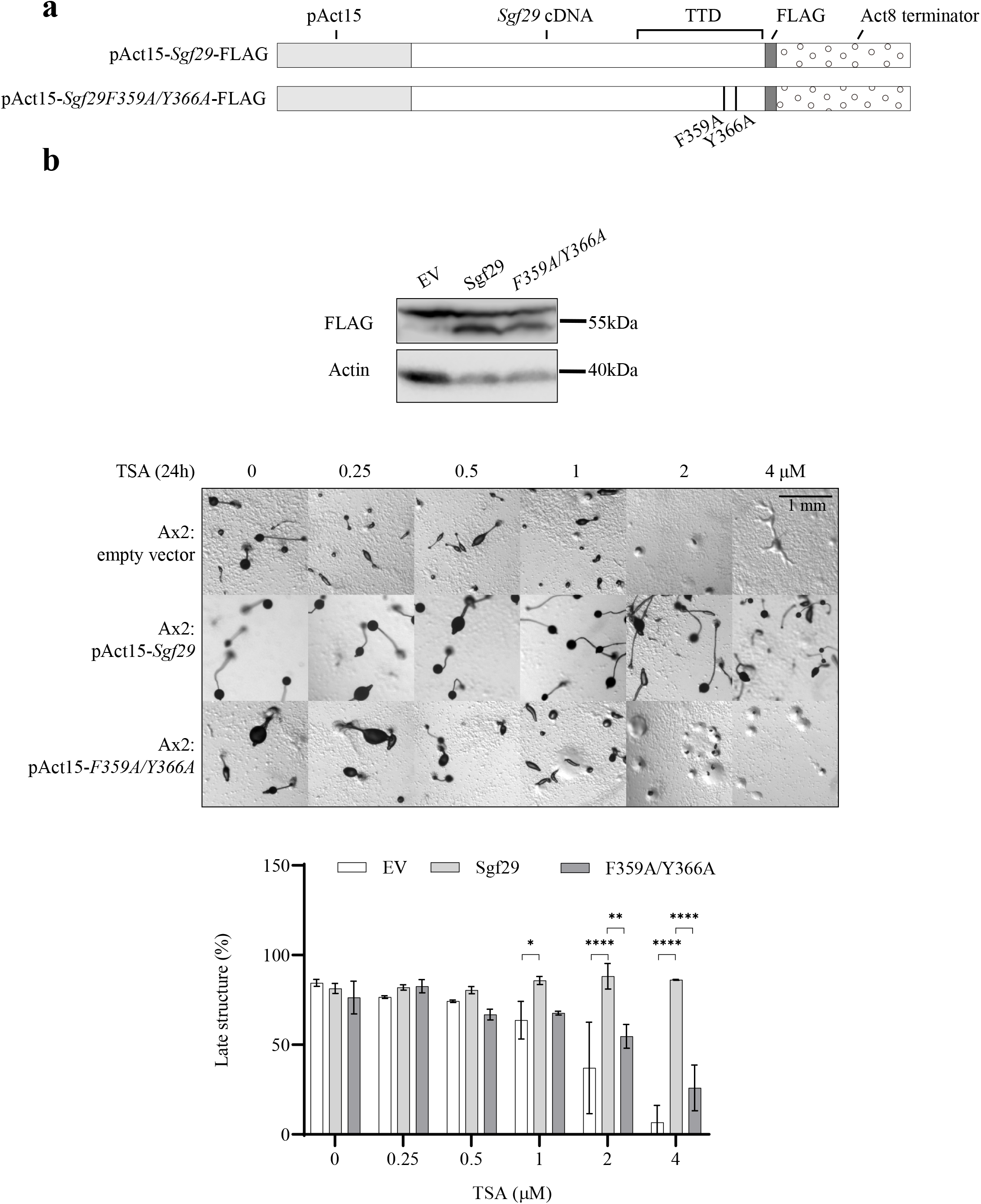
Overexpression of Sgf29-FLAG confers resistance to TSA during development. **a** Schematic illustration of wildtype and F359A/Y366A Sgf29-FLAG overexpression constructs. The expression of *C*-terminally FLAG-tagged protein is driven by the actin 15 promoter. **b** Overexpressing Sgf29-FLAG in Ax2 cells confers resistance to TSA during development. Upper panel, immunoblots showing expression of FLAG-tagged wildtype and F359A/Y366A Sgf29 in Ax2 cells. Whole cell lysates from pools of cells transfected with empty vector (EV), or constructs to drive expression of wildtype and F359A/Y366A Sgf29 were resolved by 12% SDS-PAGE and blotted using an anti-FLAG antibody. Actin is used as a loading control. Lower panel, cells were washed with KK2 and allowed to develop on solid agar with increasing concentration of TSA for 24 hours as described in Fig. 1a. Images were taken at 24 hours and representative images of three repeats were presented. Lower panel, percentage of late structures was calculated and shown as average ± SD. Statistical significance was calculated u sing two-way ANOVA with Tukey test. Statistical significance was calculated by comparing to 0h using one-way ANOVA with Dunnett test. *, *p* < 0.05; **, *p* < 0.01; ****, *p* < 0.0001.

## Discussion

Despite the use of compounds with KDACi activity in treatment of diseases such as cancer, the molecular mechanisms by which these molecules bring about long term biological effects on cells is unclear. The impact of these compounds on development is of particular interest as a number of cancer cells are induced to differentiate on therapeutic exposure to KDACi[36]. However not all cancer cells are sensitive to KDACi treatment so there is a need for biomarkers to identify those with therapeutic potential. Sensitive cancers often acquire resistance during treatment regimes, so an understanding of the mechanism of resistance is valuable to combat this. Here we exploit the sensitivity of development of *Dictyostelium* to the KDACi TSA and the relative simplicity of the histone modification machinery in *Dictyostelium* (with single copy genes encoding histone variants and reduced complexity of modifying enzymes) to explore resistance mechanisms and identify potential biomarkers. We identify the rapid, preferential acetylation on K9 and K14 of histone H3 already modified by trimethylation of K4 as an important pool of histones for mediating the sensitivity of development to KDACi.

KDACs have been implicated in the gene expression changes associated with a number of developmental processes. For example in haematopoiesis fine-tuned lineage-specific gene expression is required for the correct establishment of downstream cell types, a process that requires multiple KDACs as transcription cofactors[18]. For instance, the HDAC1-containing NuRD complex is required for the recruitment of GATA1, which promotes expression of β-globin and is essential for erythroid differentiation in mice[37–39]. Inhibition of KDACs by SAHA pre-treatment or deletion of HDAC3 in hematopoietic stem cells from mice results in loss of self-renewal, which is indicative of loss of multilineage potential[40].

In *Dictyostelium* the concentrations of TSA tested phenotypically block late development, past the aggregation stage. However short term exposure of cells during the first four hours following starvation is necessary and sufficient to confer inhibition, illustrating that the short-term effect of TSA exposure has long-term impact on *Dictyostelium* development. Although no change in overall acetylation levels of H3 on K9 or K14 are normally apparent during the first four hours of development, exposure to TSA during this time period does lead to a detectable increase in both modifications. Importantly, this acetylation is preferentially targeted to histone H3 molecules already containing the H3K4me3 modification. This anchoring of the immediate rise of H3 acetylation by to H3K4me3 histones has been reported in mouse, *Drosophila* and human cells[26] and we have previously reported that this is conserved in proliferating vegetative *Dictyostelium* cells[30]. This is the first report that this targeting is also found in cells entering differentiation.

To understand if this preferential acetylation of H3K4me3 is a relevant biological target of TSA, we investigated the developmental sensitivity of cells lacking detectable H3K4me3 firstly by deletion of the gene encoding Set1[29]. Developing *set1*^−^ cells show resistance to TSA and a delayed increase in H3 K9/14 acetylation, consistent with this pool of histones being a biologically relevant target. Set1 may have other targets so the importance of histone methylation was confirmed by replacing the endogenous genes encoding H3 with a mutated version with an alanine in this position to block methylation[30]. There are two major H3 variants expressed in *Dictyostelium*, H3a and H3b, both encoded by single copy genes facilitating the introduction of these mutations. We have previously reported that the H3aK4A gene replacement strain lacks any detectable tri-methylation of H3a or H3b, although some mono- and di-methylation of H3b is apparent[30]. Conversely, H3bK4A cells do still have detectable H3aK4me3, so these strains distinguish the role of trimethylation from mono-di-methylation, as well as importance of the two histone variants. The immediate elevation of H3 K9/14Ac in response to TSA treatment correlates with the existence of H3K4me3 as it is reduced if only mono and di-methylation are detectable. Notably, in H3bK4A cells, a TSA-induced increase in K9Ac on H3a (which still has detectable H3K4me3) is apparent, though in wild type cells the H3b variant is most highly acetylated. In H3aK4A cells, where H3bK4me1/2 are detected, no increase in acetylation of H3b is detected, suggesting that these modifications do not have the same targeting ability. Both of these strains show resistance to TSA-induced developmental inhibition, suggesting that acetylation of H3bK4me3 histones is sufficient to cause the developmental phenotype. No strain is available that only blocks methylation of H3a, so it cannot be ruled out that there is also a consequence of the targeting on this variant. In all strains there is correlation between H3bK4me3, preferential K9/14 acetylation and resistance to the developmental block induced by TSA.

It could be argued that changes in gene expression caused by loss of H3K4me3 are the root cause of developmental resistance to TSA, not the consequent preferential acetylation. Therefore, we set out to generate a strain in which the methylation is preserved but the link between H3K4me3 and H3 acetylation is broken. In *S. cerevisiae* the Sgf29 protein provides such a link as it contains a TTD that binds H3K4me3 and is associated with the GCN5 HAT complex. The protein coded by DDB_G0272054 is a strong candidate for the *Dictyostelium* orthologue of Sgf29 as it is the only protein encoded in the genome with a TTD. This TTD interacts *in vitro* specifically with H3K4me3 and not with histones lacking this mark extracted from *set1^−^*, H3aK4A and H3bK4b cells and this binding is dependent on residues known to be required for this interaction in Sgf29[33]. The binding of recombinant protein to H3b in H3aK4A cells is undetectable, even though H3K4me1/2 is still detectable on H3b in these cells [30]. The *C*-terminal TTD of both human and budding yeast Sgf29 favours binding to H3 peptides containing H3K4me3 over H3K4me2 although both interactions are detectable[33]. H3K4me3 has been shown to be the preferred ligand and Sgf29 is localised at gene promoters by ChIP-seq analysis, overlapping with the H3K4me3 modification[41]. Any TTD binding to H3K4me2 histones is below the detectable limit by our technique. One factor contributing to the limitation could be that there is around half the level of H3b compared to H3a reducing the amount of H3b with H3K4me2 to be detected by the recombinant protein[30].

Disruption of the *sgf29* gene is not lethal, in line with studies in other organisms including *S. cerevisae* and *C. neoformans*[33, 42] and is not required for *Dictyostelium* development. In *C. neoformans*, disruption of *sgf29* leads to hypervirulence, and inability to form titan cells, and in Arabidopsis with salt tolerance, demonstrating important biological consequences of its loss[42–44]. Importantly disruption of the *sgf29* gene successfully disrupts preferential H3K9/14 acetylation of H3K4me3 upon TSA treatment and also confers resistance to TSA during development. We cannot rule out changes in levels of H3K4me3 at individual gene promoters as has been reported for ER stress response genes in human cells[45]. The basal level of H3K9 and K14 acetylation in *sgf29*^−^ cells is similar to control cells (Fig 4D) which differs from the situation in *S. cerevisiae* where loss of Sgf29 leads to a reduction in global acetylation levels[33] even though the structural integrity of the SAGA complex[46]. However, delay of H3K9Ac and H3bK14Ac accumulation on TSA treatment of developing cells was found in *sgf29*^−^ cells, which indicates a general delay of histone H3 acetylation upon TSA treatment and in particular the kinetics of acetylation of H3K4me3 is not distinguishable from that of bulk H3. This agrees with a previous finding that loss of the TTD in *S. pombe* Sgf29 results in loss of acetylation to H3K4me3 nucleosomes *in vitro* but the reactivity of HAT module remains the same[47].

Overall our data is consistent with the identified protein being the orthologue of Sgf29 and that in *Dictyostelium* early development, it is responsible for the preferential H3K9/14 acetylation of H3K4me3 in the presence of TSA and this, in turn, plays a major role in the sensitivity of development to TSA. The TSA-resistance is dependent on the presence of TTD. Mutation of the TTD at residues F359 and Y366 was sufficient to lose *in vitro* binding activity but cells expressing full length protein with the equivalent mutation show an intermediate resistance phenotype, suggesting some residual binding activity *in vivo* or another chromatin targeting domain in the complex. This is consistent with *S.cerevisiae* Sgf29 where mutation of individual binding residues reduced H3 acetylation but not to the level of complete deletion of Sgf29[33]. Overexpressing Sgf29 in Ax2 cells also dramatically enhances resistance to TSA. One possibility is that free excess protein binds to H3K4me3 and blocks recruitment of the acetyltransferase complex. This raises the interesting possibility that either loss or overexpression of Sgf29 in tumour cells can act as a therapeutic biomarker for potential resistance to hydroxamates such as TSA. In the COSMIC database (https://cancer.sanger.ac.uk/cosmic), over 10% of tested patient samples of breast, haematological, large intestine and stomach cancers show overexpression of Sgf29 as well as 5 out of 8 cancer cell lines, compared to non-transformed lines[48]. This overexpression it has been proposed to lead to upregulation of expression of the oncogene c-myc, for example in hepatocellular carcinomas[49]. It will be informative to investigate whether these cells show altered sensitivity to hydroxamates and whether other components of this pathway such as reduced levels of H3K4me3 seen in some cancers, could also provide biomarkers for resistance [50, 51].

## Conclusions

This work reveals the importance of preferential acetylation of histone H3 proteins that are previously modified by trimethylation of lysine 4 in the mode of action of the histone deacetylase inhibitor TSA in inhibition of development. *Dictyostelium* strains that lack this methylation, either through loss of the Set1 enzyme that deposits it, or through mutation of the endogenous histone H3 genes to replace the lysine with an alanine, are resistant to the inhibitory effects of TSA on development. To demonstrate that the effect is not due to loss of methylation directly, we generated a strain in which methylation is conserved but lacks a protein, Sgf29, that recognises the modification. Loss of Sgf29 prevented preferential acetylation of this pool of methylated histones and also led to resistance to TSA in development. This is the first description of a biological role for preferential acetylation of this pool of histone in the mechanism of action of this class of inhibitor. As this family of inhibitors is used to treat some cancers, this knowledge will be important in identifying cancers which are sensitive to this class of drugs.

## Methods

### *Dictyostelium* growth and development

*Dictyostelium discoideum* axenic strain Ax2 cells were grown axenically in HL5 media in shaking suspension at 220 rpm at 22° C. Exponentially growing cells were used for all experiments. The development assay was carried out in a 24-well plate format. 0.5 ml 1.5% agar (Formedium) in KK2 (19 mM KH_2_PO_4_, 3.6 mM K_2_HPO_4_) were added to each well and allowed to set and dry in a laminar flow hood using maximum air flow for 2h with lid open. DMSO vehicle control or TSA was applied on top of the agar to the desired concentration and the plates dried. Exponentially growing cells were collected, washed three times with KK2 then resuspended to 1.4 × 10^8^ cells/ml in KK2 and 20 μl applied to each well and dried for 30 mins. Cells were allowed to develop for 24 hours. Pictures were taken using a camera (Hamamatsu, model ORCA-05G) attached to a dissection microscope (Leica, model MZFLLIII). The number of early and late structures were manually counted. In TSA pre-treatment experiments, exponentially growing cells were collected, washed and resuspend to 4 × 10^6^ cells/ml in KK2 supplemented with 4 μM TSA and shaken at 120rpm at 22 degrees. Cells were the plated and allowed to develop on solid agar as mentioned above after 4 hours treatment or subjected to acid extraction after 0, 1, 2 or 4 hours treatment in shaking suspension.

### Generation of *sgf29*^−^ cells

A plasmid was constructed to disrupt the *sgf29* gene by homologous recombination. The 5’ arm (1 to 1085 of *sgf29*) was amplified and cloned into pJET1.2 vector followed by insertion of amplified 3’ arm (1943 to 2877) at XhoI site. Next, the bsR cassette was obtained from the pLPBLP vector by SmaI digestion and subsequently inserted at the HincII site between the 5’arm and 3’arm, generating the final disruption vector pJET_Sgf29KO. Sequences of primers used and the map of the final construct are included in additional files 6 and 7. The strategy replaces the coding sequence from 1086 to 1942 with a blasticidin resistance cassette (BsR), disrupting expression downstream of amino acid 119, whereas the TTD starts at aa 277. The disruption vector was digested with BsrGI and BsyZ17I and electroporated into Ax2 cells. Following selection with 10 μg/ml blasticidin, individual resistant colonies were screened for disruption of the *sgf29* gene by PCR (Additional File 4: Figure S4). Three independent clones were used to verify the developmental resistance and loss of H3K4me3-directed acetylation in response to TSA.

### Cloning and purification of GST-TTD from *E. coli* cells

The sequence coding for the tandem Tudor domain (TTD) (S277 - K411) in Sgf29 (411 a.a.) was amplified from genomic DNA. Mutation of the TTD (F359A/Y366A) was generated by two stages of PCR. Two individual fragments containing a single mutation were firstly amplified and used as templates for a second stage of amplification. Wildtype or mutant Tudor domain was then subcloned into the pGEX-KG vector using EcoRI (5’) and XhoI (3’) sites. This generated an expression cassette with GST fused to the N-terminus of the TTD and the expression in *E. coli* was driven by an IPTG-inducible (Isopropyl β-d-1-thiogalactopyranoside, Sigma) promoter.

Transformed *E.coli* were inoculated into 50 ml LB containing 100 mg/ml ampicillin (LB-Amp) and cultured at 37° C O.N. The starting culture was diluted into 600 ml LB-Amp the next day at 37° C until O.D 600 reached 0.55. One mL of the culture was stored before IPTG was added to a final concentration of 0.1 mM. The culture was then incubated at 16° C O.N. for maximum protein expression. One mL of the O.N. culture was stored before cells were collected and lysed by sonication (10 watts, 10s intervals with 1-minute cooling in between) in 15 ml lysis buffer (50 mM Tris, 50 mM NaCl and 1X protease inhibitor cocktail from Roche) at 4° C. Cell lysates were centrifuged at 48,000 × g for 20 min at 4° C to separate soluble protein in supernatant from inclusion bodies in the precipitates, both of which were stored for SDS-PAGE analysis. The GST-TTD fusion protein was purified using Glutathione sepharose 4B (GE Healthcare) following the manufacturer’s instructions. Samples collected from various stages mentioned above, including flow through sample and purified GST-TTD fusion proteins, were resolved by 10% SDS-PAGE to follow the purification process and the purity of the final product. Sequences of primers and the final constructs used are listed in Additional file 6 and 7.

### Cloning and overexpression of full-length *Sgf29* in *Dictyostelium*

Full length wildtype and F359A/Y366A Sgf29 were cloned by two steps. First, the coding sequence of 1 to 854 of *Sgf29* was amplified from cDNA. The N-terminal end of the amplicon carried a BamHI cutting site introduced by the forward primer while the C-terminal end of the fragment overlapped with the previously cloned wildtype and F359A/Y366A TTD. The N-terminal fragment was cloned into pGEX-KG carrying wildtype or F359A/Y366A TTD using BamHI and NcoI sites, generating full-length Sgf29. The full-length Sgf29 cassette also carries BglII and SpeI sites at 5’ and 3’ ends, respectively. The two sites allowed subcloning of the full-length Sgf29 cassette into the pDM304 vector containing *C*-terminal 1xFLAG tag and allows protein expression in *Dictyostelium* driven by an Actin15 promoter [52]. Sequences of primers and the final constructs used are listed in Additional file 6 and 7.

### Expression of*Sgf29* driven by endogenous promoter in *Dictyostelium*

The endogenous promoter of *sgf29* (pSgf29, 824 bp upstream of start codon) was amplified and cloned into pDM304 vectors containing wildtype or F359A/Y366A Sgf29 with a *C*-terminal 3xFLAG tag using XhoI and BglII sites to replace the original Actin15 promoter. The expression cassette (pSgf29-Sgf29-3xFLAG) was then subcloned into pJET_Sgf29KO to replace the 5’homology arm using BtgI site. The BsR cassette in *sgf29*-null cells was removed by cre-lox recombination as previously described[35]. The rescuing vector was digested using BsrGI and BsyZ17I and electroporated into cre-loxed *sgf29* null cells. Cells surviving blasticidin selection (10 μg/ml) were pooled and evaluated for expression of 3xFLAG-tagged Sgf29 and subjected to development assays.

### Acid extraction for histone enrichment

A total of 2 × 10^8^ exponentially growing cells were pelleted at 1,700 × g for 3 min, washed in 25 ml of cold KK2 twice and lysed using 2 ml of AE lysis buffer (50 mM Tris pH 8.0, 20 mM NaCl, 3 mM MgCl_2_, 3 mM CaCl_2_, 0.5 M Sorbital 0.6% Triton X-100, 10 mM Sodium butyrate supplemented with 25 μl Phosphatase Inhibitor Cocktail 2&3 (P5726&P0044, Sigma) and 1 tablet of proteinase inhibitor cocktail (A32963, Thermo Fisher) in 5 ml lysis buffer). Nuclei were collected by centrifugation at 2,500 × g at 4 °C and washed twice by 1 ml of AE wash buffer (lysis buffer without salts and supplemented with 100 mM β-mercaptoethanol) at 4°C. Nuclei were then subjected to acid extraction using 250 μl 0.4M HCl for 1h at 4°C. Thee sample was centrifuged at 16, 000 × g for 15 min to separate supernatant and insoluble proteins. The supernatant was transferred to a new 2 ml tube and the proteins were precipitated by adding 6.2X volume of cold acetone. Samples for SDS-PAGE and AU gel were separated at this step with one fourth (SDS-PAGE) of the total sample transferred into one Eppendorf and the rest sample (AU gel) remained in the original tube. Proteins are allowed to precipitate O.N. at 4°C. Samples were washed twice with 1 ml cold acetone after O.N. precipitation.

### Protein electrophoresis and western blot

The preparation of different percentages of SDS-PAGE and performing of electrophoresis were carried out as described in “Molecular Cloning” (Green and Sambrook, 2012). Acid-Urea gel electrophoresis was carried out as previously described [26]. Acid extracts were resolved by 18% SDS-PAGE or 20% Acid-urea gel then transferred to a PVDF membrane. Antibodies against H3 (1:3000, Abcam #12079), H3K9Ac (1:3000, [26, 30]), H3bK14Ac (1:3000), H3K4me3 (1:10000,[26, 30]), β-actin (1:3000, Santa Cruz Biotechnology #sc-47778), FLAG (1:3000; Sigma, #F3165). Odyssey Fc imaging system (LI-COR, Lincoln) was used to image blots. Rabbit anti-H3bK14Ac was generated using peptide SSQ-K(Ac)-SFPSTQGLC-KLH.

## Declarations

### Ethics Approval and Consent to participate

Not applicable

### Consent for publication

Not applicable.

### Availability of data and material

All data generated or analysed during this study are included in this published article and its additional information files.

### Author Contributions

L-Y.H and C.P designed the study. L-Y.H. and D-W.H performed the experiments, L-Y.H. and C.P. analysed and interpreted the data and wrote the manuscript.

## Acknowledgments

We would like to thank Pia Theissen for technical support on antibody characterisation.

---

**Fig S1.**
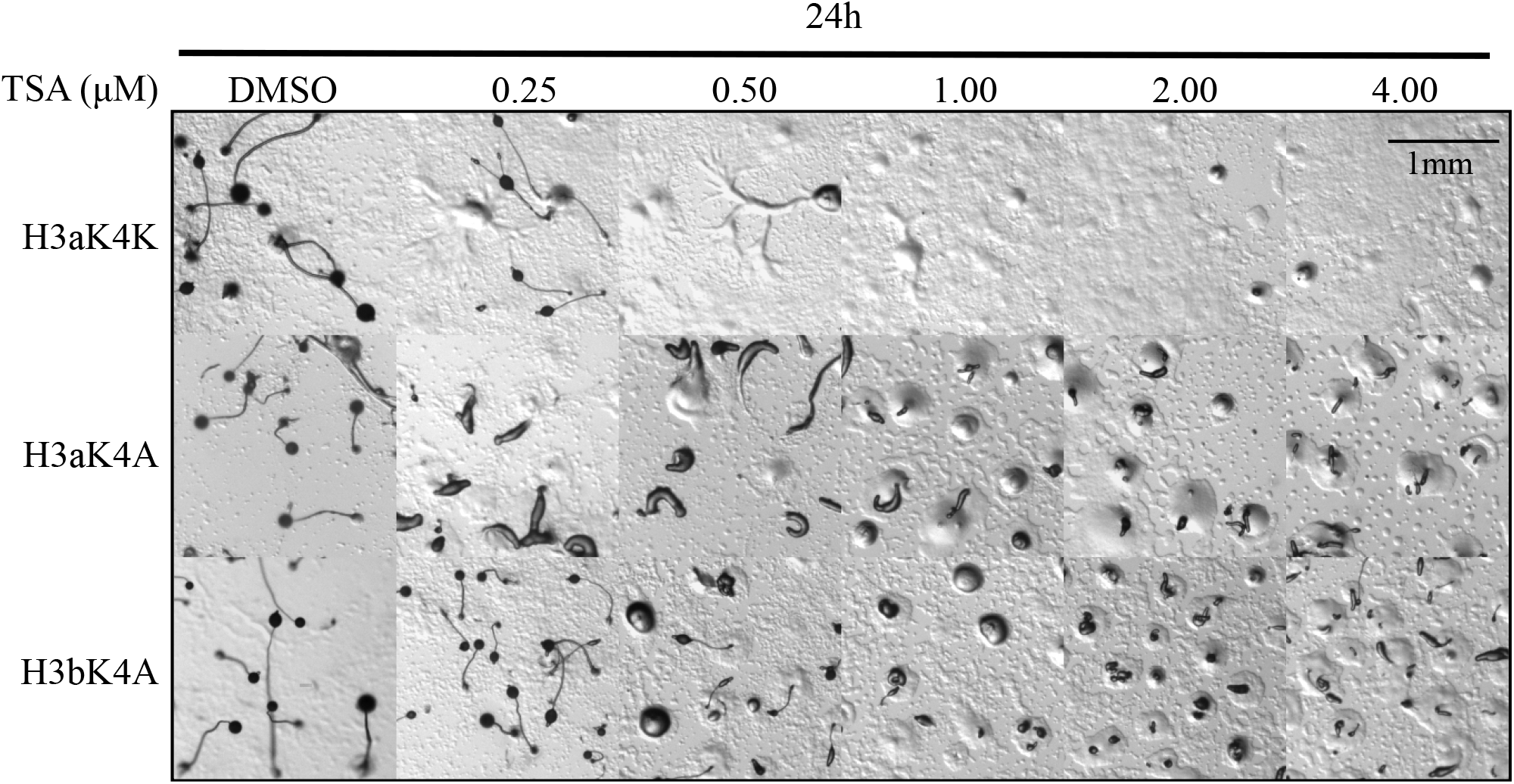
Development of H3aK4A and H3bK4A cells after 24 hours of TSA treatment Exponentially growing H3aK4A, H3bK4A or control H3aK4K cells were collected and washed twice with KK2 then transferred to 1.5% agar (1.5 × 10^6^ cells/cm^2^) containing increasing concentrations of TSA or DMSO vehicle control. Images were taken after 24 hours of development. Scale bar represents 1 mm.

**Fig S2.**
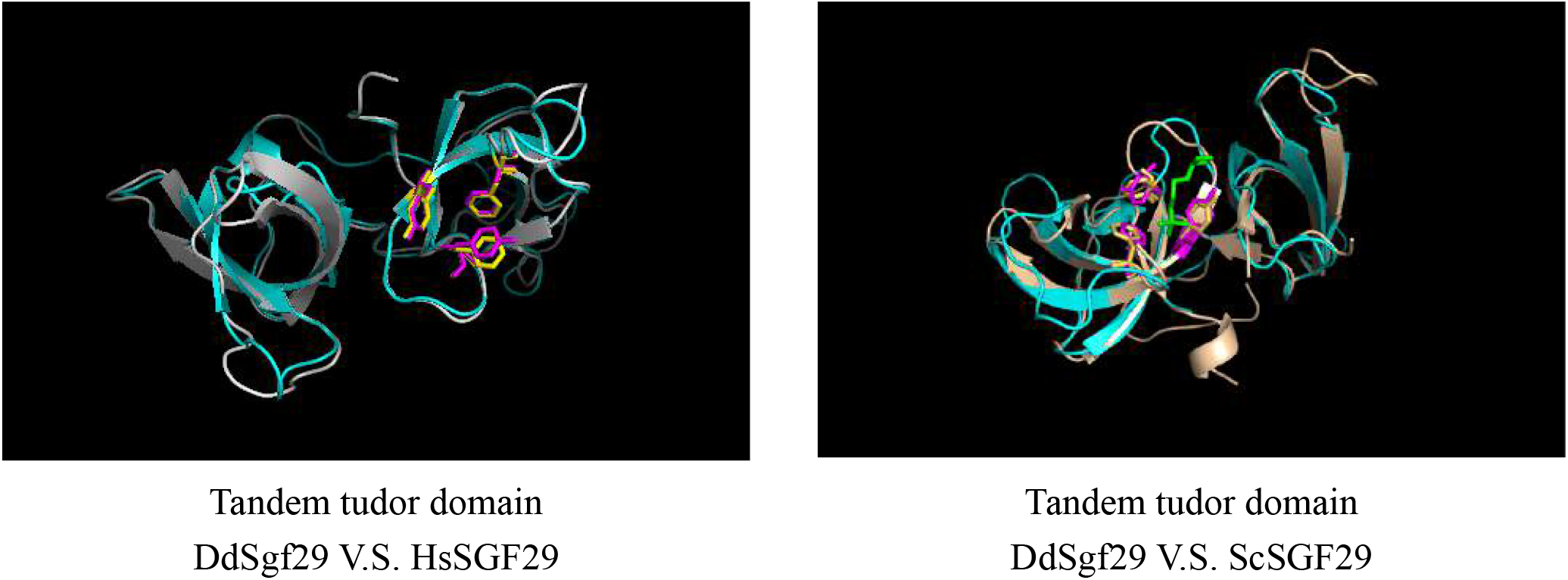
Structure alignment of the TTD of *Dictyostelium* Sgf29 to budding yeast and human counterparts. The predicted tertiary structure of the TTD of *Dictyostelium* Sgf29 is shown by a blue ribbon while that of human (HsSGF29) or budding yeast (ScSGF29) is shown in grey. Conserved binding residues in *Dictyostelium* Sgf29 are shown as purple sticks. K4-trimethylated lysine is represented as a green stick. Structural prediction and comparison were done using Chimera (https://www.cgl.ucsf.edu/chimera/).

**Fig S3.**
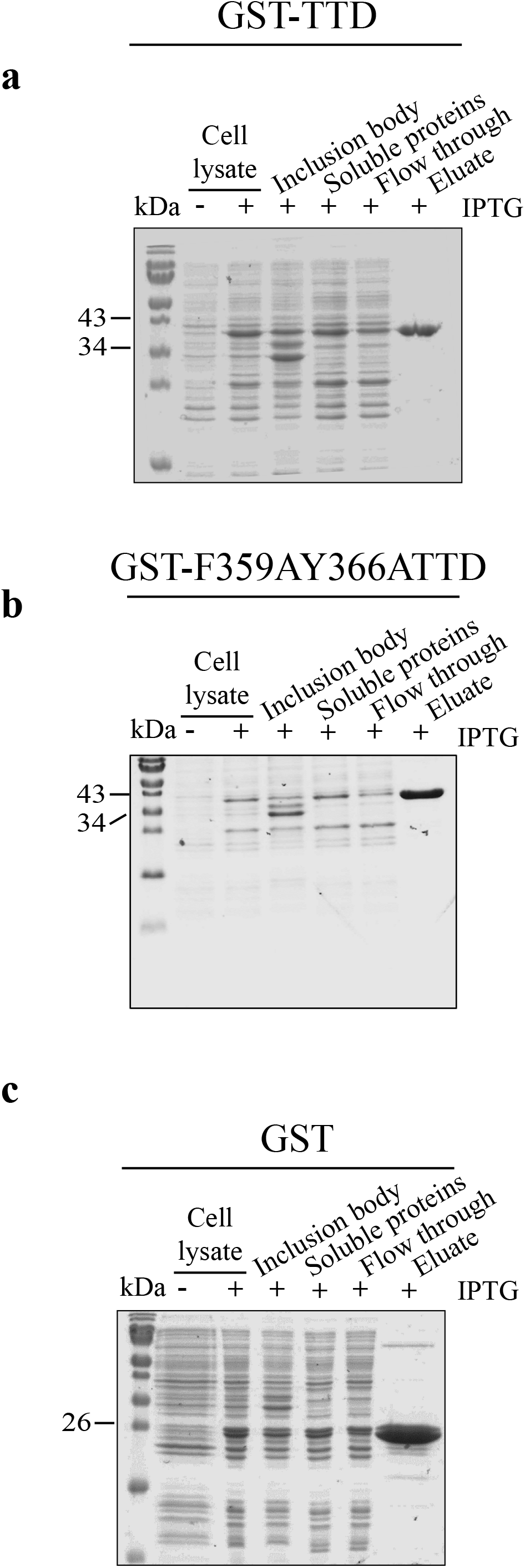
Purification of GST-TTD fusion proteins following expression in *E.coli*. GST-fusion proteins were purified using glutathione sepharose. Samples from stages of the purification were resolved by SDS-PAGE and proteins were visualized by staining with Coomassie blue for **a** GST-TTD (42 kDa), **b** GST-F359AY366ATTD (42 kDa) and **c** GST (29kDa) in individual gels.

**Fig S4.**
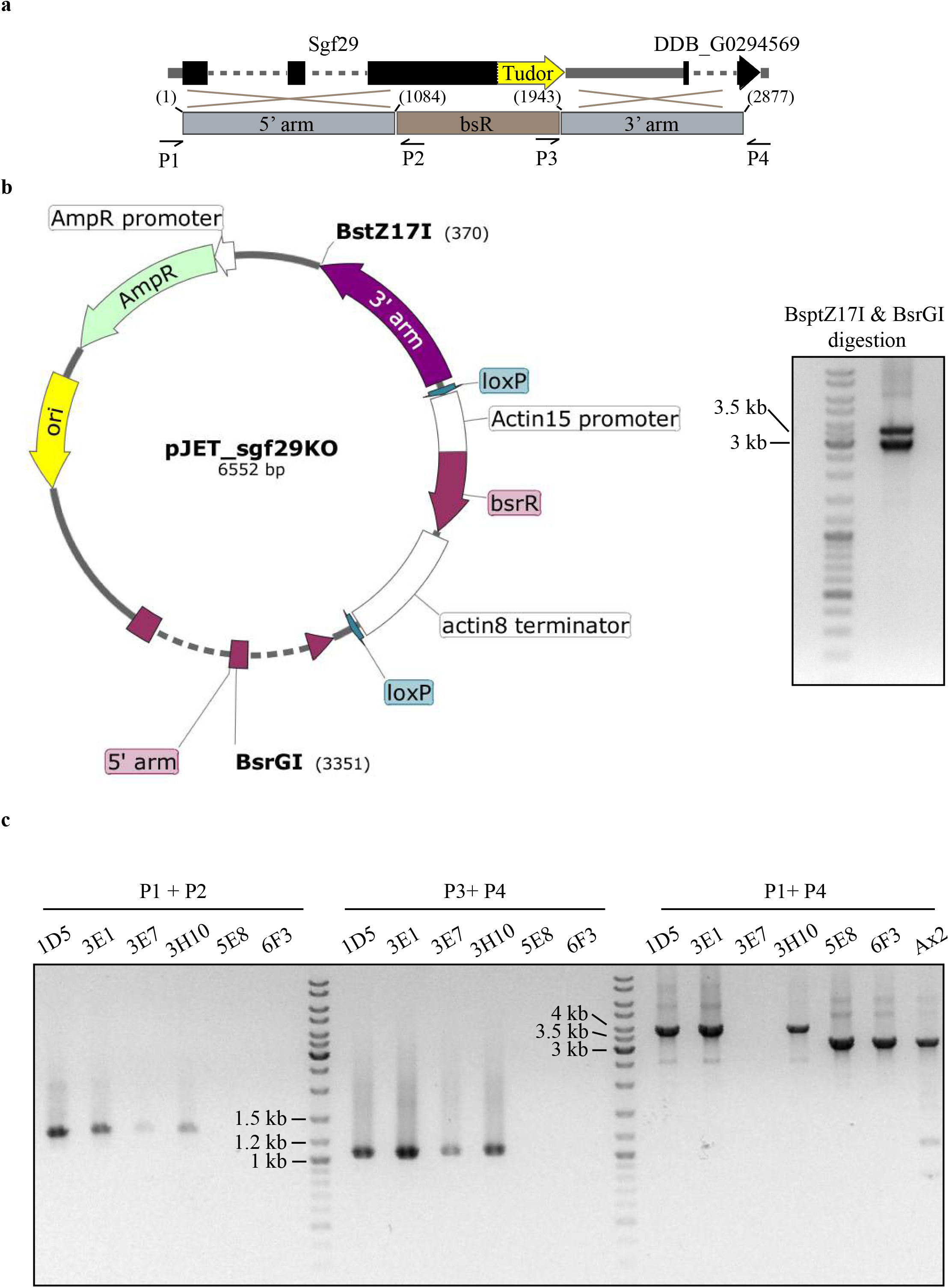
Generation and genotyping of *sgf29*^−^ cells **a** Schematic representation of the gene replacement event to generate *sgf29*^−^ cells. Exon, intron and intergenic regions are represented by a thick black solid line, a dashed line and a solid grey line respectively. The TTD, marked yellow, is located at the 3’end of the coding region. The 5’ arm spanning 1 to 1084 relative to the start codon, 3’ arm (1943 to 2877) and the BsR cassette was cloned into pJET1.2 vector to generate pJET_sgf29KO disruption vector. Specific primer sets were designed to detect bsR insertion (P1, & P2; P3 & P4). A primer set (P2 & P4) flanking the linearized fragment was used to detect parental contamination. **b** Map of the pJET_sgf29KO disruption vector. The vector was digested using BsrGI and BstZ17I before transfected into parental Ax2 cells, as shown on the agarose gel. **c** gDNA was prepared from surviving clones after blasticidin selection. PCR screening was carried out using primer sets mentioned above. Gene replacement at the correct sites were detected by primer P1 and P2 with the PCR product at 1368 bp while that of P3 and P4 was at 1091 bp. Parental signal was detected using primer set P2 and P4, with a band at 3121 bp while the size after gene replacement is 3854 bp. Clone 1D5 (clone 1), 3E1 (clone 2) and 3H10 (clone 3) are disruptants and clone 1 is used for main figures.

**Fig S5.**
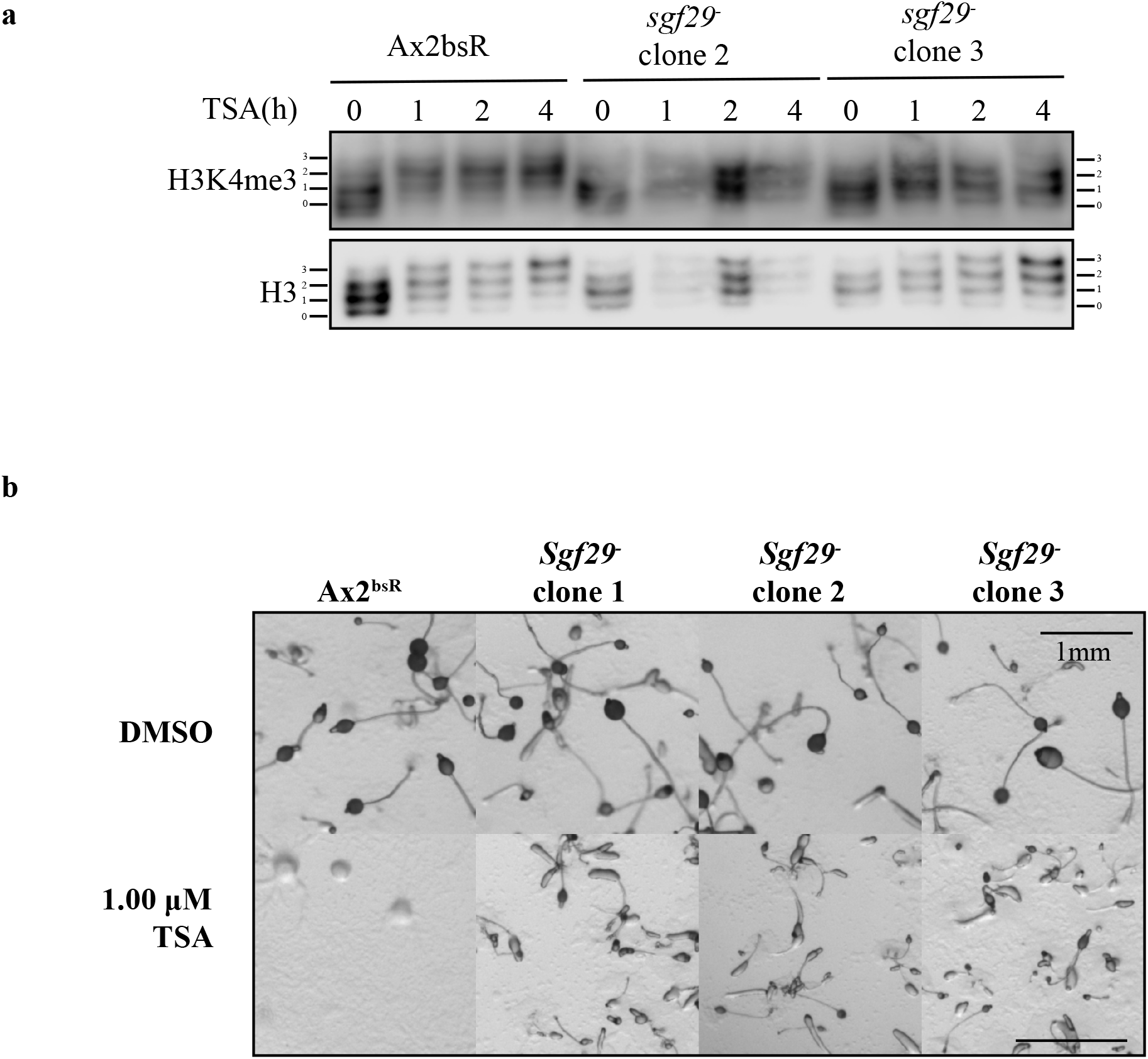
Verification of *sgf29*^−^ phenotypes using independent clones. **a** Histone-enriched acid extracts from TSA (4 μM) treated developing Ax2^bsR^ and two further independent clones of *sgf29*^−^ cells were prepared after 0, 1, 2 or 4 hours of TSA treatment. Samples were resolved by 20% acid-urea electrophoresis and western blots were performed using anti-H3K4me3 and anti-H3 polyclonal antibodies. Position of histone H3 with decreasing net positive charge (0 – 3) are shown on both sides of each blot. **b** Cells from three independent *sgf29*^−^ clones and Ax2^bsR^ controls were washed with KK2 and allowed to develop on solid agar with 1μM TSA for 24 hours as described in Fig. 1a. Images were taken at 24 hours and representative images of three repeats are presented. Clone 1 is used for main figures.

## Additional Files

Additional File 1: Figure S1.pdf

Additional File 2: Figure S2.pdf

Additional File 3: Figure S3.pdf

Additional File 4: Figure S4.pdf

Additional File 5: Figure S5.pdf

Additional File 6: Primers.

Additional File 7: Plasmid maps.PDF

## Additional File 6_Primers

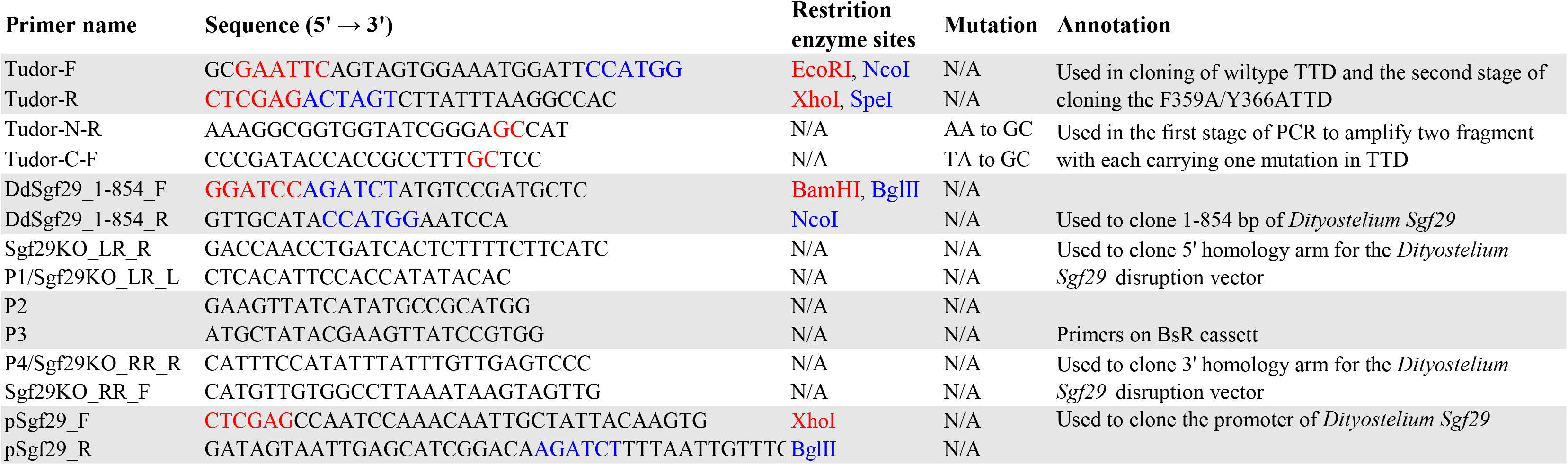

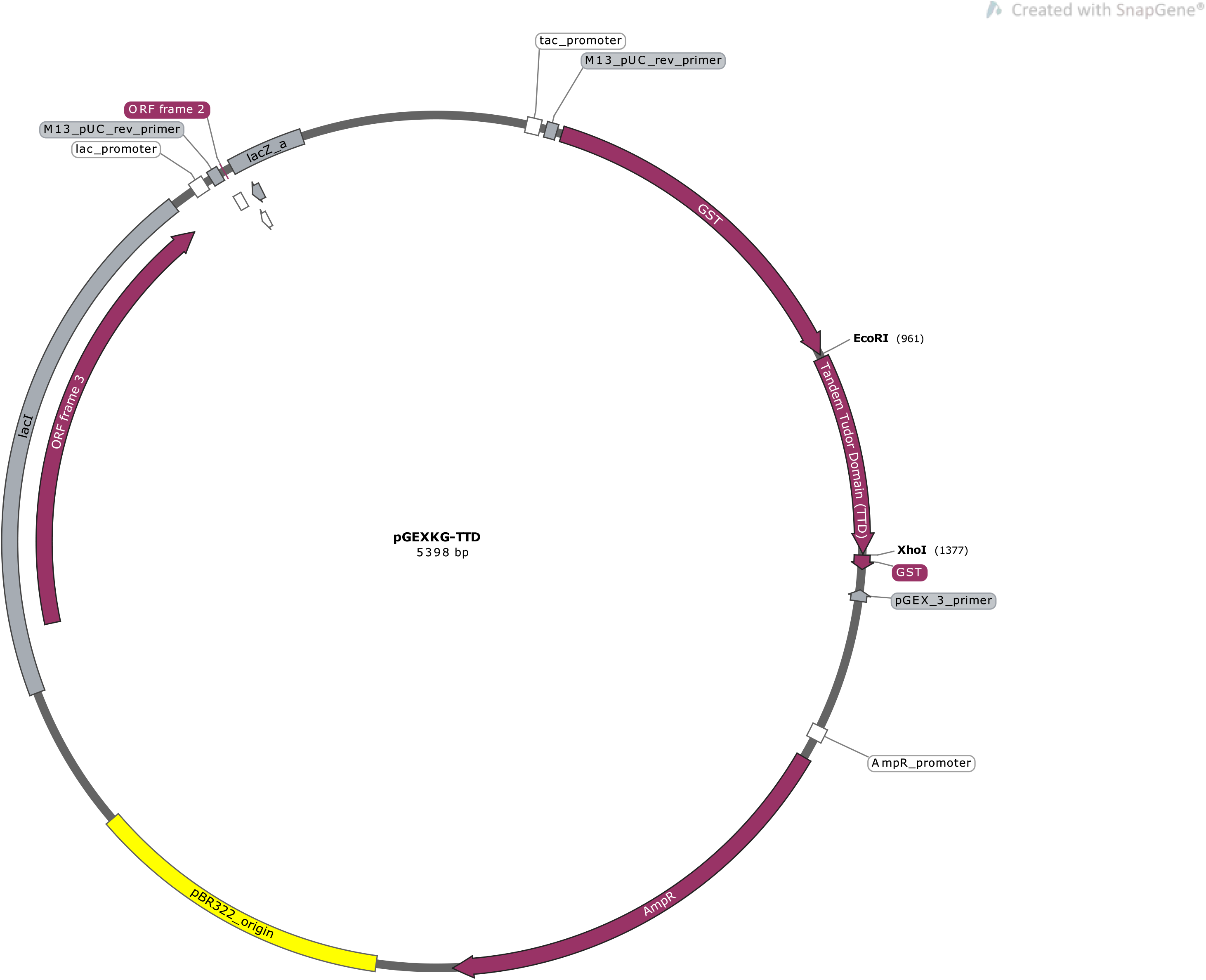

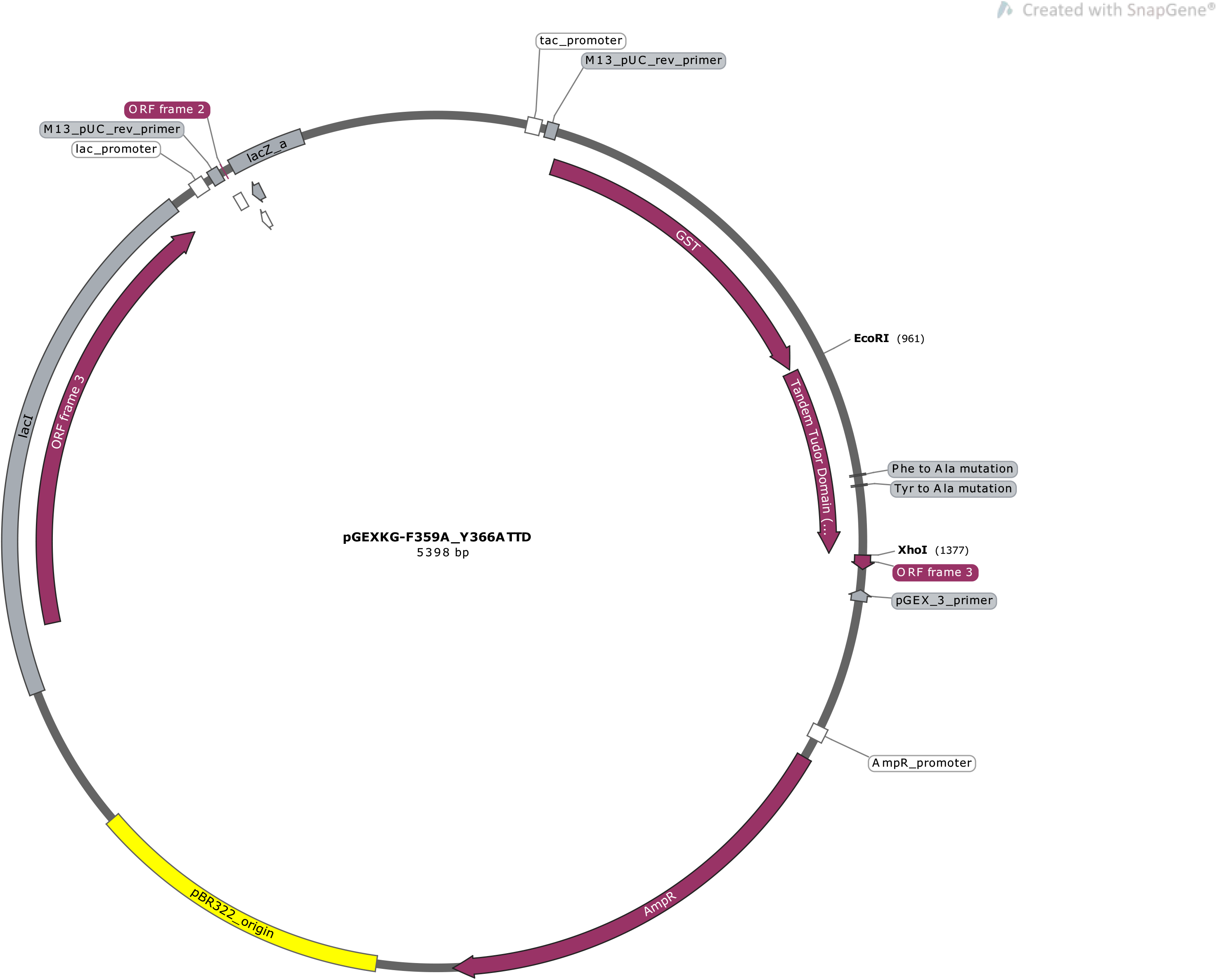

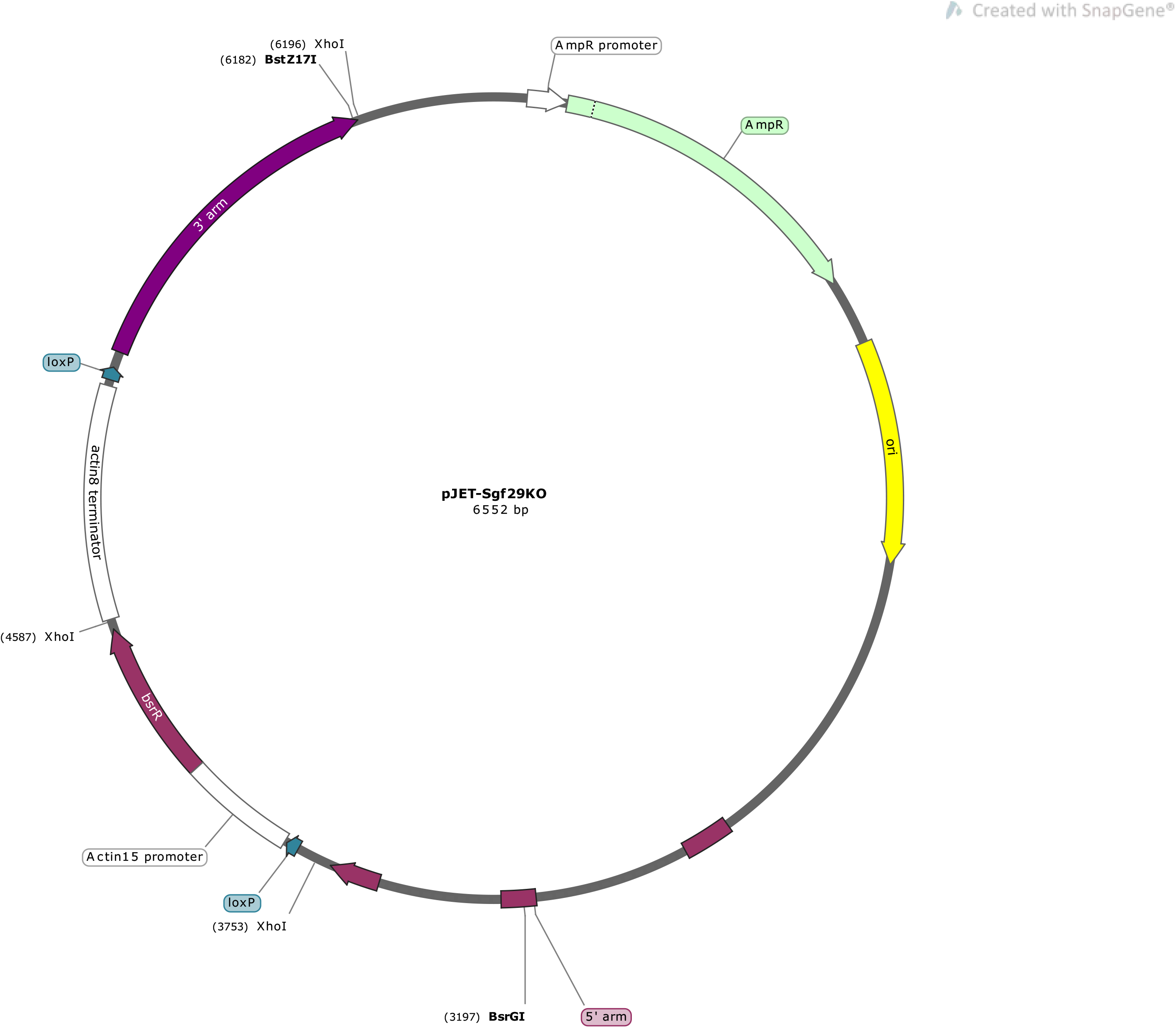

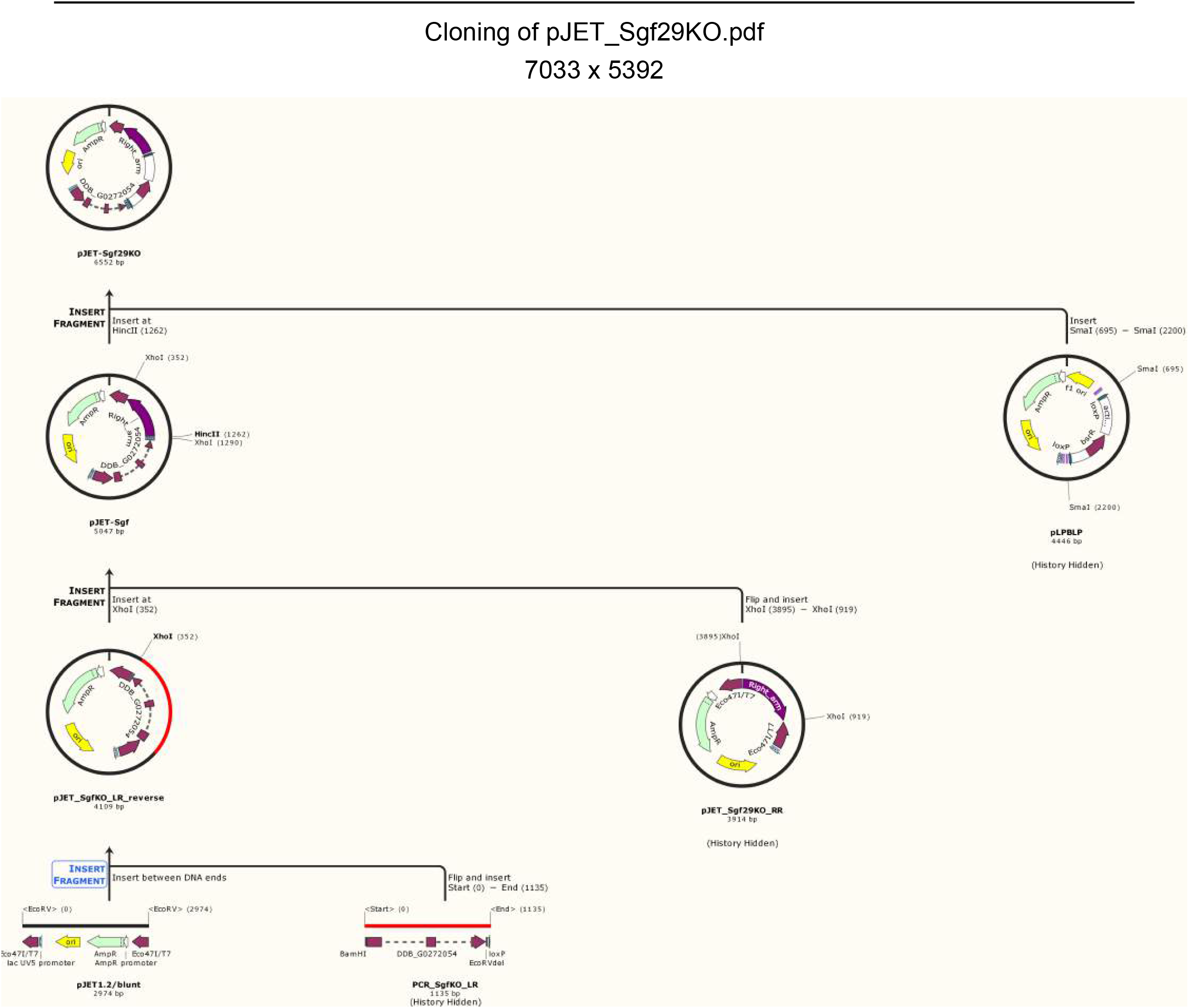

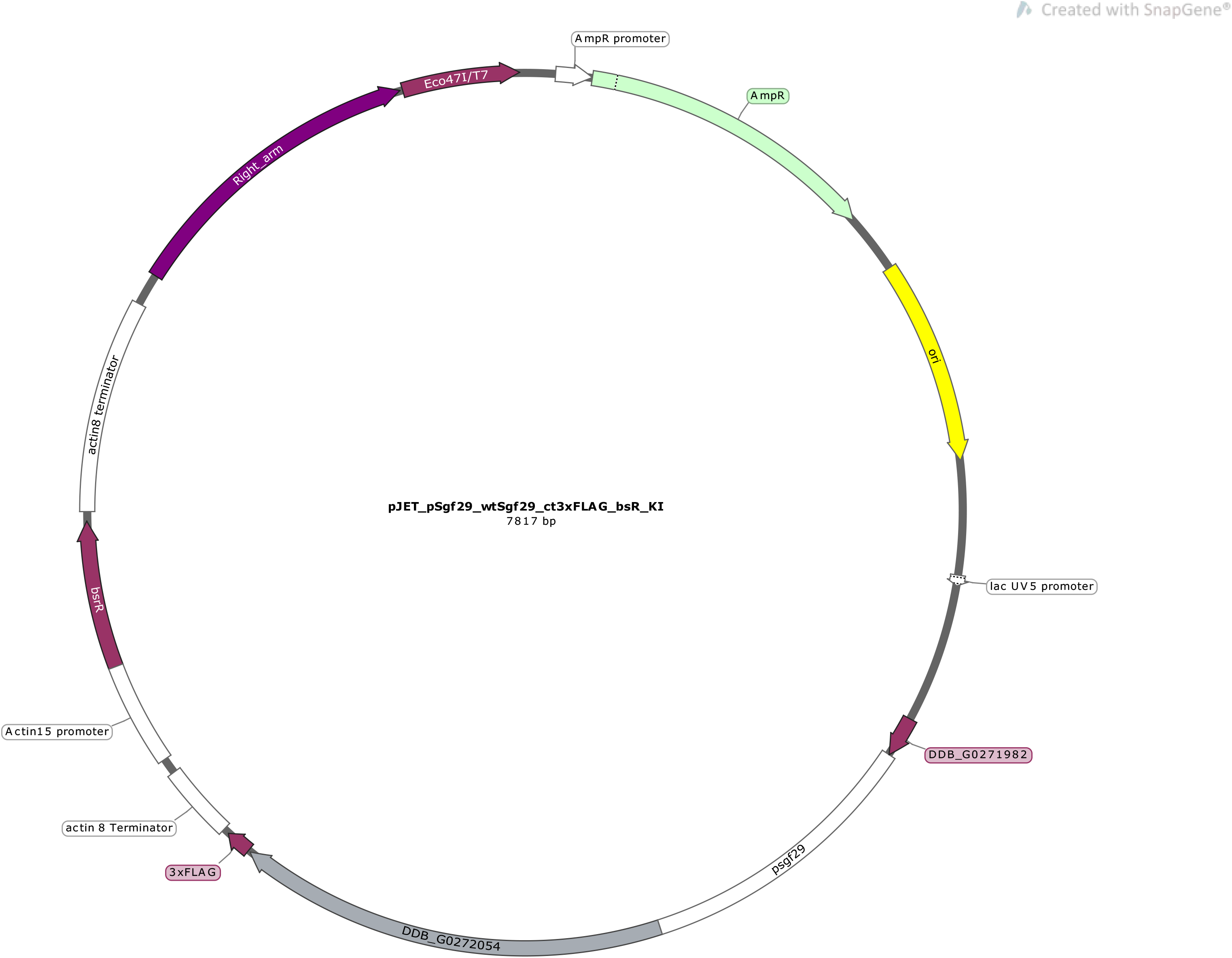

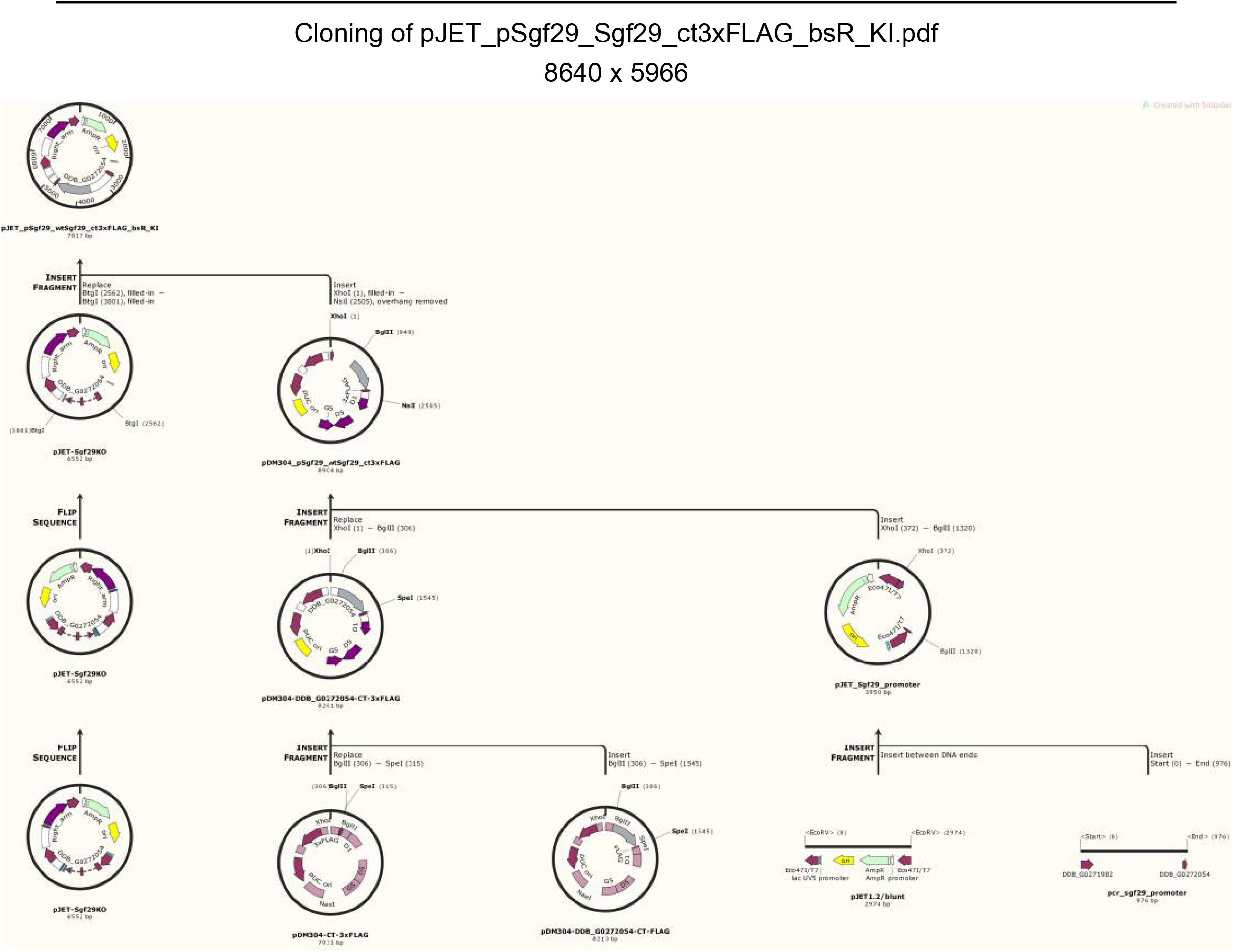

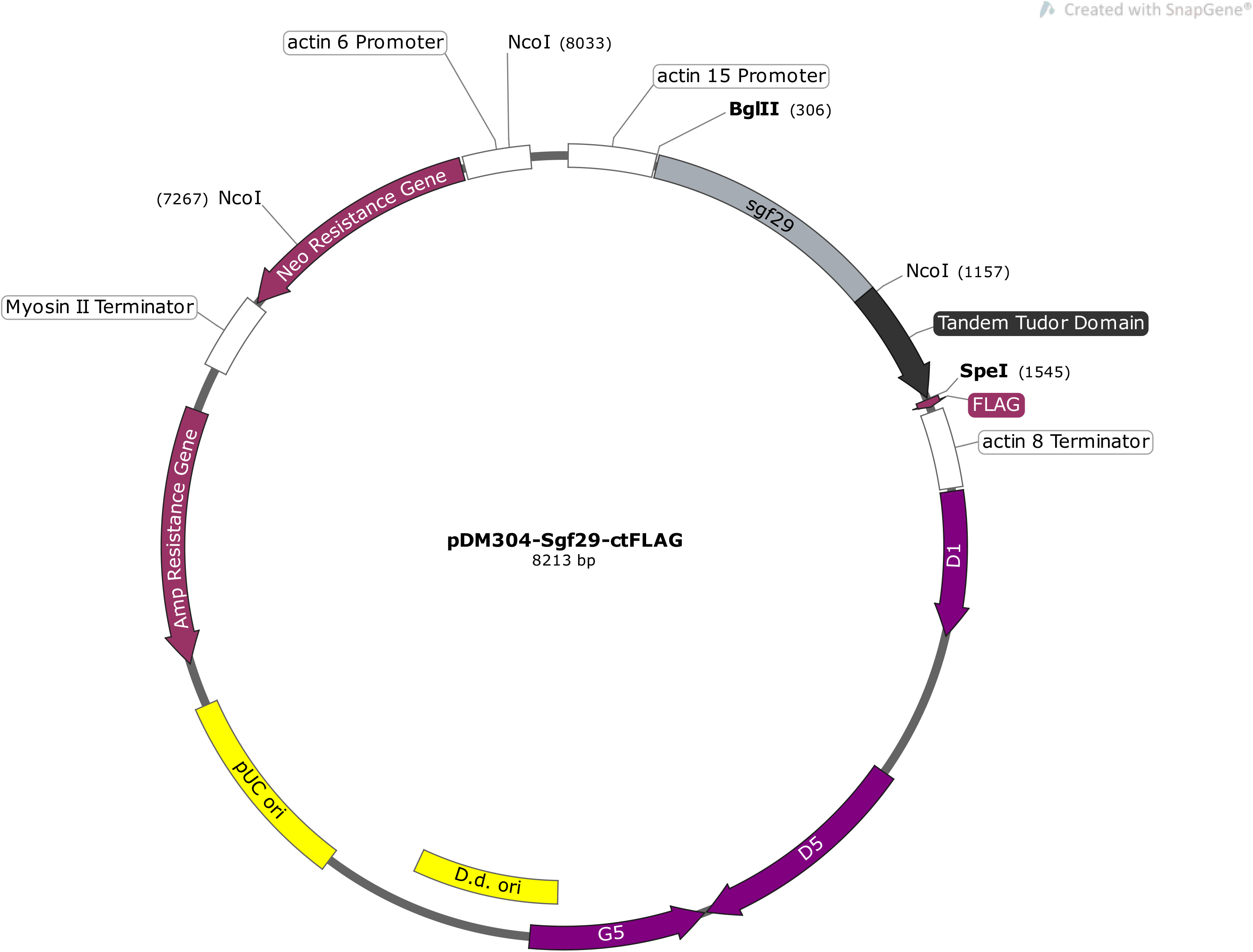

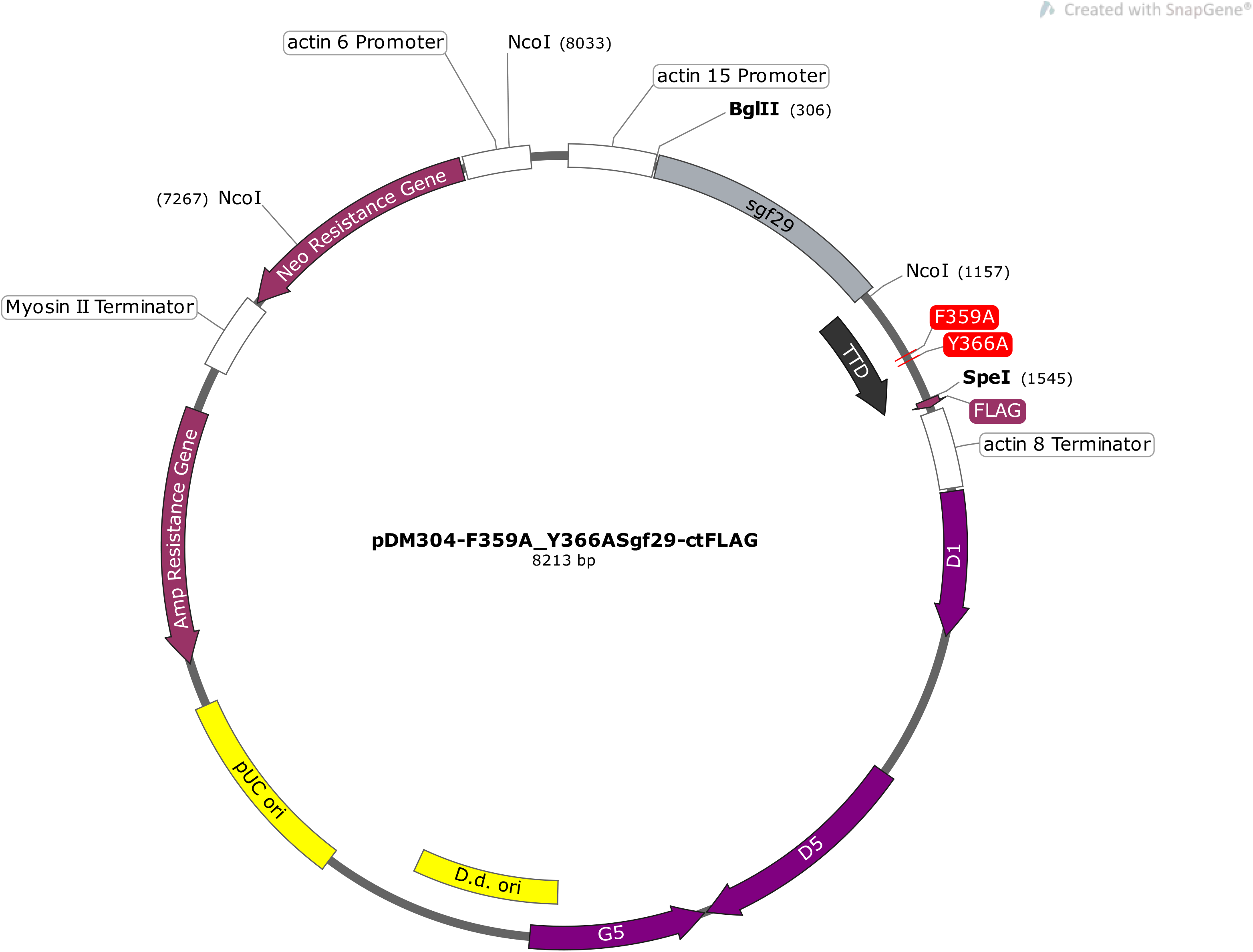

